# Tau assemblies enter the cytosol in a cholesterol sensitive process essential to seeded aggregation

**DOI:** 10.1101/2021.06.21.449238

**Authors:** Benjamin J. Tuck, Lauren V. C. Miller, Emma L. Wilson, Taxiarchis Katsinelos, Shi Cheng, Marina Vaysburd, Claire Knox, Lucy Tredgett, Emmanouil Metzakopian, Leo C. James, William A. McEwan

## Abstract

Accumulating evidence supports a prion-like mechanism in the spread of assembled tau in neurodegenerative diseases. Prion-like spread is proposed to require the transit of tau assemblies to the interior of neurons in order to seed aggregation of native, cytosolic tau. This process is poorly understood and remains largely hypothetical. Here, we develop sensitive techniques to quantify the cytosolic entry of tau in real-time. We find that tau does not promote its own entry but, rather, is wholly dependent on cellular machinery. We find that entry to the widely used reporter cell line HEK293 requires clathrin whereas entry to neurons does not. Cholesterol depletion or knockdown of cholesterol transport protein Niemann-Pick type C1 in neurons renders cells highly vulnerable to cytosolic entry and seeded aggregation. Our findings establish entry as the rate-limiting step in seeded aggregation and demonstrate that dysregulated cholesterol, a feature of several neurodegenerative diseases, potentiates tau aggregation.

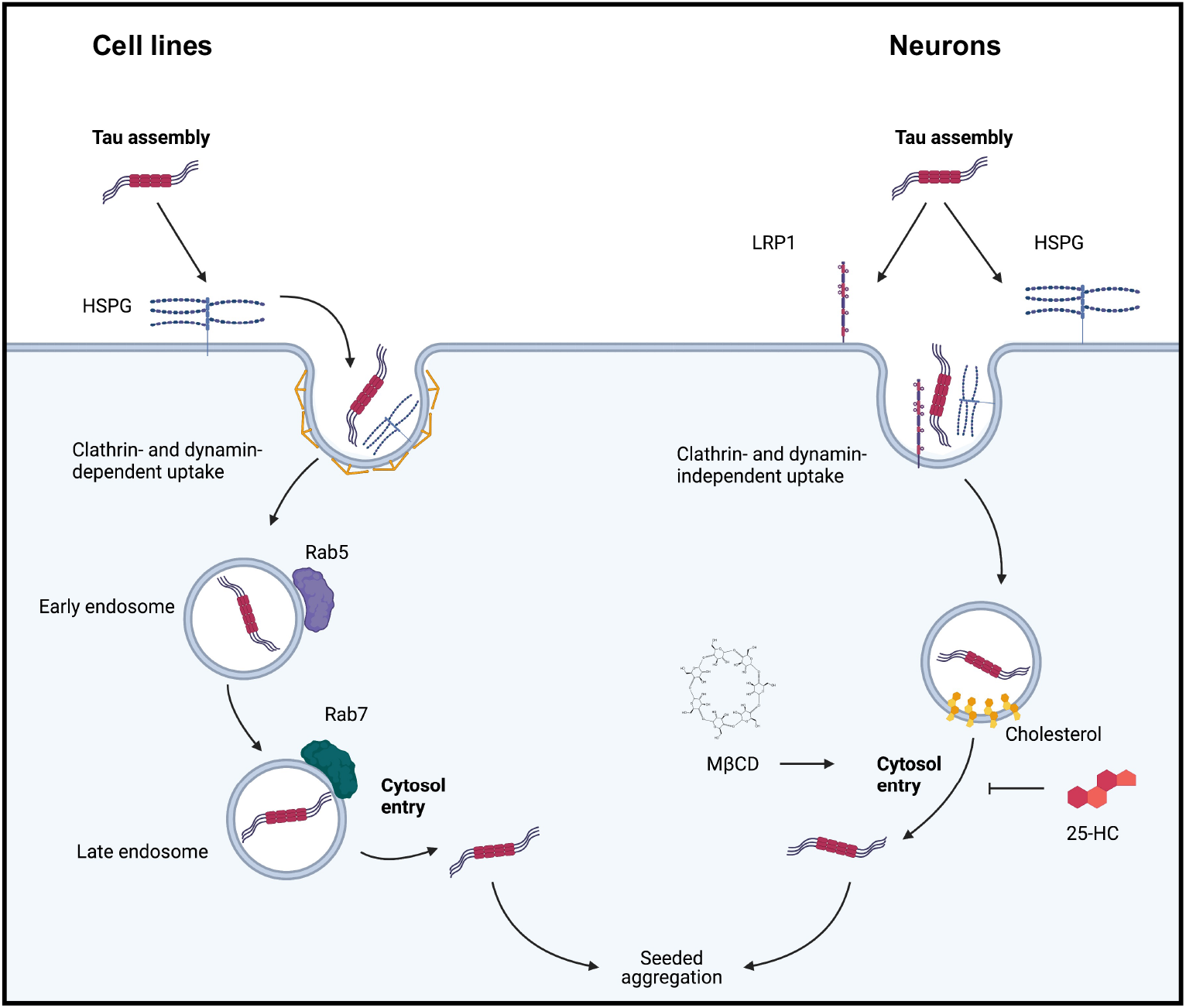
Graphical Abstract.

## Introduction

Tauopathies are a group of neurodegenerative diseases characterised by the conversion of microtubule-associated protein tau into highly ordered fibrils^1^. Most prominent among these diseases is Alzheimer’s disease, which accounts for the majority of the ∼50 million dementia cases worldwide. Other tauopathies include fronto-temporal dementia, progressive supranuclear palsy and chronic traumatic encephalopathy. A causal role for tau in neurodegeneration is indicated by more than 50 nonsynonymous and intronic point mutations which lead to dominantly inherited, early onset forms of neurodegenerative disease characterized by the accumulation of tau assemblies in the brain^2^. Two non-mutually exclusive mechanisms are proposed to explain the occurrence of tau fibrils in the diseased brain. First, that nucleation of aggregation occurs within individual cells in a cell autonomous manner. Alternatively, that tau assemblies transit between cells, promoting the aggregation of tau in recipient cells in a ‘prion-like’ manner. This latter model of aggregation could help explain the observed spatio-temporal spread of tau misfolding in the human brain and is consistent with observations of seeded aggregation in cultured cells^3–6^, and in *in vivo* disease models^7–10^. Nonetheless, the relative contributions of these two mechanisms to human disease progression remains unknown^11^. Spread between cells may occur through the uptake of naked tau assemblies, or via extracellular membrane-bound vesicles. Naked tau assemblies are taken up into membrane bound vesicles following interactions between tau and cell-surface heparan sulphate proteoglycans (HSPGs), as well as the recently identified low-density lipoprotein receptor, LRP1^12,13^. Tau is then taken up to membrane-bound compartments via endocytosis and macropinocytosis^12,14–16^. For a prion-like mechanism to occur, tau assemblies must subsequently gain access to the cytosol, somehow breeching these intracellular membranes. While the process of tau filament uptake has been comparatively well studied, the transfer of tau from intracellular membrane-bound vesicles to the cytosol is largely unexplored and remains a critical missing step for assessing the relevance of seeded aggregation to disease progression^11,17^.

Cholesterol is a critical determinant of membrane bilayer structural integrity, and a known risk factor in neurodegenerative disease^18–21^. Cholesterol is depleted from the brain in an age-dependent manner, resulting in impaired intracellular signaling and synaptic plasticity^22–24^. Variations in *APOE*, which encodes a cholesterol transporting protein, is the strongest genetic risk factor for AD with inheritance of the ε4 variant being associated with a substantially increased risk of AD^25,26^. Though a link between APOE and Abeta pathology is well established, experimental evidence also suggests that APOE exacerbates tau pathology independently of Abeta^27–30^. Supporting a role of cholesterol biology in tau pathology, *APOE* was also identified as a risk factor in primary age-related tauopathies, where Abeta is not implicated^31^. In the primary tauopathies, the ε2 allele, which is protective against AD, is associated with higher risk. Niemann-Pick type C (NPC) is a rare autosomal recessive disorder characterised by the aberrant accumulation of intracellular cholesterol and glycolipids. Approximately 95% of NPC cases are caused by loss-of-function mutations in the Niemann-Pick C1 gene (*NPC1*), which encodes a ubiquitous trafficking protein important for transport of cholesterol to organelles and the plasma membrane^32,33^. NPC is characterized by progressive childhood neurological disease often with abundant tau pathology. Thus, cholesterol abundance and localization are strongly associated with tau pathology and may play a direct role in its progression.

In this study we develop highly sensitive methods for the detection of tau entry to the cytosol, permitting analysis of entry at physiological concentrations of tau. We find that entry of tau to the cytosol represents the rate-limiting step in seeded aggregation. Accordingly, changing entry levels through pharmacological or genetic means results in a concomitant change in seeded aggregation. We find that the entry pathway of tau assemblies into human embryonic kidney cells (HEK293), a widely used reporter line, differs substantially to entry in mouse or human neurons. We observe that tau does not mediate its own entry to the cytosol of neurons but, rather, enters through a clathrin- and dynamin independent pathway. Tau entry to neurons relies on cell surface receptor LPR1, heparin sulphate proteoglycans and membrane cholesterol. Depletion of cholesterol from the plasma membrane, or its mislocalisation by NPC1 depletion, enhances tau entry in multiple cell models, including organotypic slice culture and human neurons. The results establish tau entry as a distinct event from uptake that is essential for seeded aggregation. They further demonstrate that tau entry is modulated by cellular determinants rather than by tau itself and is therefore potentially susceptible to pharmacological inhibition.

## Results

### Recombinant Tau-HiBiT assemblies can reconstitute NanoLuc *in vitro* and intracellularly

The study of tau entry to cells has been complicated by the difficulty in reliably distinguishing cytosolic populations from vesicular populations^17^. In order to specifically detect the cytosolic fraction of exogenously supplied tau assemblies, we established a live-cell assay relying on split luciferase, NanoLuc (Nluc) (**Fig. 1a**). NanoLuc is a 19 kDa luminescent protein engineered from the luciferase of a deep-sea shrimp *Oplophorus gracilirostris.* The enzyme has been split into an 18 kDa subunit (LgBiT) of Nluc and an 11 amino acid peptide (HiBiT) which interact with sub nanomolar affinity. Reconstitution results in the complementation of activity and luminescence in the presence of substrate^34^. We expressed recombinant P301S tau (0N4R isoform) in fusion with a HiBiT tag at the C-terminus in *E. coli* (**Fig. 1b,c**). Assemblies of tau-HiBiT were produced by incubation with heparin and the aggregation kinetics were quantified via thioflavin T fluorescence, a dye which fluoresces when bound to β-sheet amyloid structures^35^. We observed similar aggregation kinetics compared to a tagless variant of tau, suggesting that the HiBiT tag does not interfere with aggregation (**Fig. 1d**). Negative stain electron microscopy revealed filamentous structures with smaller assemblies present after sonication (**Fig. 1e**). To assess the ability of tau-HiBiT assemblies to reconstitute Nluc, we titrated tau-HiBiT assemblies into a fixed concentration of recombinant LgBiT, resulting in a concentration-dependent increase in luminescent signal upon the addition of substrate (**Fig. 1f**). We found the HiBiT tag was as readily accessible in assemblies as it was in monomer, as signal was unchanged between these states (**Fig. S1a**). These results confirm that tau-HiBiT can form filaments similar to tagless variants and these assemblies yield a dose-dependent luminescent signal when complexed with LgBiT.

**Figure 1:**
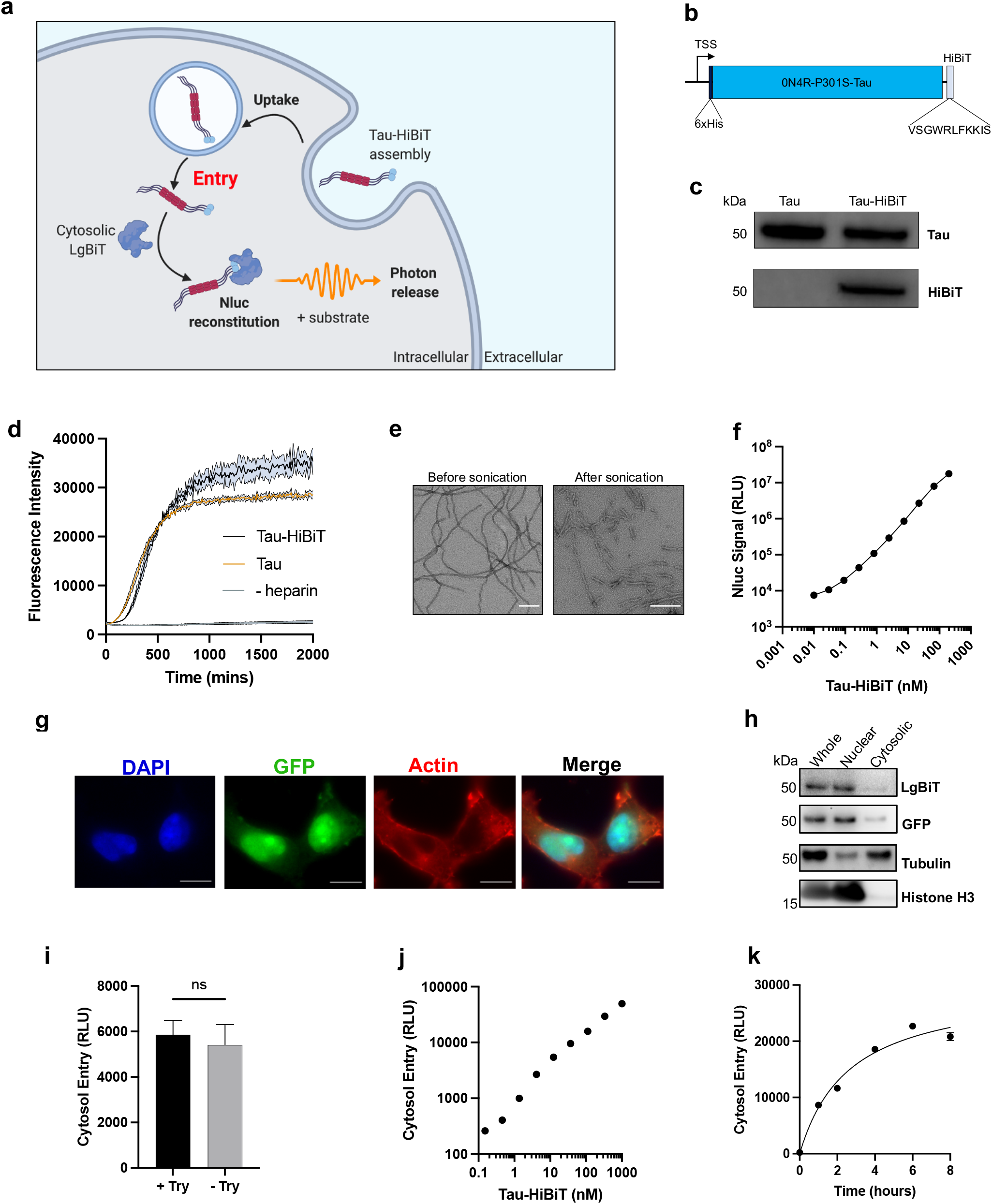
Characterisation of HiBiT-tagged tau assemblies and their entry to the cytosol of HEK293 cells. **a)** Cartoon depicting the intracellular reconstitution of NanoLuc and the enzymatic production of light through interaction of tau-HiBiT with cytosolic LgBiT. **b)** Depiction of the 6xHis-0N4R-P301S-Tau-HiBiT construct and the amino acid sequence of the HiBiT peptide, to scale relative to tau. **c)** Western blot of 50 ng recombinant tau or tau-HiBiT monomers with anti-tau (DAKO) or anti-HiBiT antibody. **d)** Time-course of 5 µM tau-HiBiT and tagless tau aggregation kinetics monitored by thioflavin T (15 µM) fluorescence; n=4. **e)** Representative transmission electron micrographs of heparin-induced tau-HiBiT assemblies before and after a single 15 second sonication cycle. Scale bar, 200 nm. **f)** Representative titration curve of tau-HiBiT assemblies complexed with recombinant LgBiT *in vitro* for 30 minutes; n=4. **g)** Immunofluorescence imaging of HEK293 cells expressing nls-eGFP-LgBiT (HEK-NGL), demonstrating nuclear compartmentalisation of NGL. Scale bars, 50 µm. **h)** Western blotting of cytosolic and nuclear fractions of NGL lysates probing for LgBiT, GFP, nuclear marker histone H3 and cytosolic marker, tubulin**. I)** Effect of 10 minutes trypsin protease treatment on luminescent signal in NGL cells prior to substrate addition and signal acquisition; n=≥3. **j)** Representative entry titration curve of tau-HiBiT assemblies added exogenously to HEK-NGL cells for 1 h; n=3. **k)** Representative time-dependency entry curve of 50 nM tau-HiBiT assemblies on NGL cells over the course of 8 h; n=3. TSS; transcriptional start site, NGL; nls-eGFP-LgBiT, nls; nuclear localisation signal, RLU; relative light units, Try; trypsin. Error bars are mean ± s.e.m.

In order to template aggregation, tau assemblies are proposed to cross cell limiting membranes. To measure the entry of tau assemblies to the cell, we expressed LgBiT in the cytosol of human embryonic kidney HEK293 cells, a cell line widely used in the study of seeded tau aggregation^3,6,36,37^. We expressed LgBiT by lentiviral transduction from the vector pSMPP ^38^ which drives expression from a viral (spleen focus forming virus; SFFV) promoter. We applied sonicated tau-HiBiT assemblies to the cell exterior. This resulted in high luminescence signal that was sensitive to the addition of trypsin, a protease which completely abrogates the luciferase signal of cell free tau-HiBiT:LgBiT complexes (**Fig. S1b,c**). We interpret this as an indication that the bulk of the interaction was extracellular, likely due to leakage or secretion of LgBiT to the media.

We therefore examined if low cytosolic LgBiT concentrations were necessary to establish an assay that reports solely on intracellular tau entry. We reduced cytosolic concentrations of LgBiT first by replacing the SFFV promoter of pSMPP with a weak mammalian housekeeping promoter^39^ (PGK) to generate the vector pPMPP. The cytosolic concentration was further reduced by expressing the protein in fusion with a nuclear localisation signal (nls) and eGFP. The resulting construct (nls-eGFP-LgBiT, abbreviated as NGL) was expressed at ∼85:15 nuclear:cytoplasmic ratio and yielded a luciferase signal that was trypsin-insensitive following treatment of cells with tau-HiBiT and addition of a cell-penetrant luciferase substrate (**Fig. 1g-i**). We next titrated tau-HiBiT onto HEK293 cells expressing the construct (HEK-NGL) and observed dose and time-dependent uptake that saturated at around 6 h (**Fig. 1j,k**). These results demonstrate that the assay reports on the real-time entry of tau to live cells with a broad and linear dynamic range.

### Tau-HiBiT assemblies enter cell lines via clathrin-mediated endocytosis

Tau seeding can be dramatically increased by the use of transfection reagents^12^, likely owing to enhanced delivery of tau assemblies to the cell interior. To test whether transfection reagents result in increased tau entry to the cytosol, we titrated tau-HiBiT assemblies in the presence and absence of Lipofectamine 2000 (LF) onto HEK-NGL cells. We observed a 50- to-100-fold increase in entry of tau to the cytosol when supplemented with LF (**Fig. 2a**). In HEK293 cells expressing P301S tau-venus^6^, we observed an attendant increase in seeded aggregation (**Fig. 2b-d**). These data demonstrate that LF promotes entry to the cell and implicates breeching of intracellular membranes as rate-limiting to seeded aggregation. We next tested whether entry could be inhibited by reducing uptake to the cell. Previous studies have shown that tau uptake occurs via endocytosis and relies on the vesicle coat protein clathrin, as well as the GTPase dynamin^14,40–42^. We pre-treated cells with the clathrin inhibitor PitStop 2 or the dynamin inhibitor Dyngo 4a. This reduced the uptake of fluorescently labelled transferrin to the cell (**Fig. S1c**) and resulted in a stark reduction in tau entry to the cytosol (**Fig. 2e**). We also tested the routinely used Na^+^/H^+^ import channel inhibitor, dimethyl-amiloride (DMA)^43,44^, which inhibits macropinosome establishment, but observed no change in entry to the HEK-NGL cells (**Fig. 2e**). Incubation of cells at 4°C, a temperature non-permissive to endocytosis, also prevented tau entry to the cell (**Fig. 2f**). Heparan sulphate proteoglycans (HSPGs) mediate the uptake of tau to intracellular compartments and promote seeded aggregation ^12,45^. Pre-treatment with surfen hydrate, an agent that binds heparan sulphate, strongly diminished entry in our assay (**Fig. 2g**) as did addition of exogenous heparin to the cell media (**Fig. S1d**). These data together confirm that HSPGs promote entry of tau assemblies to the cytosol of HEK293 cells, and that entry relies upon clathrin and dynamin.

**Figure 2:**
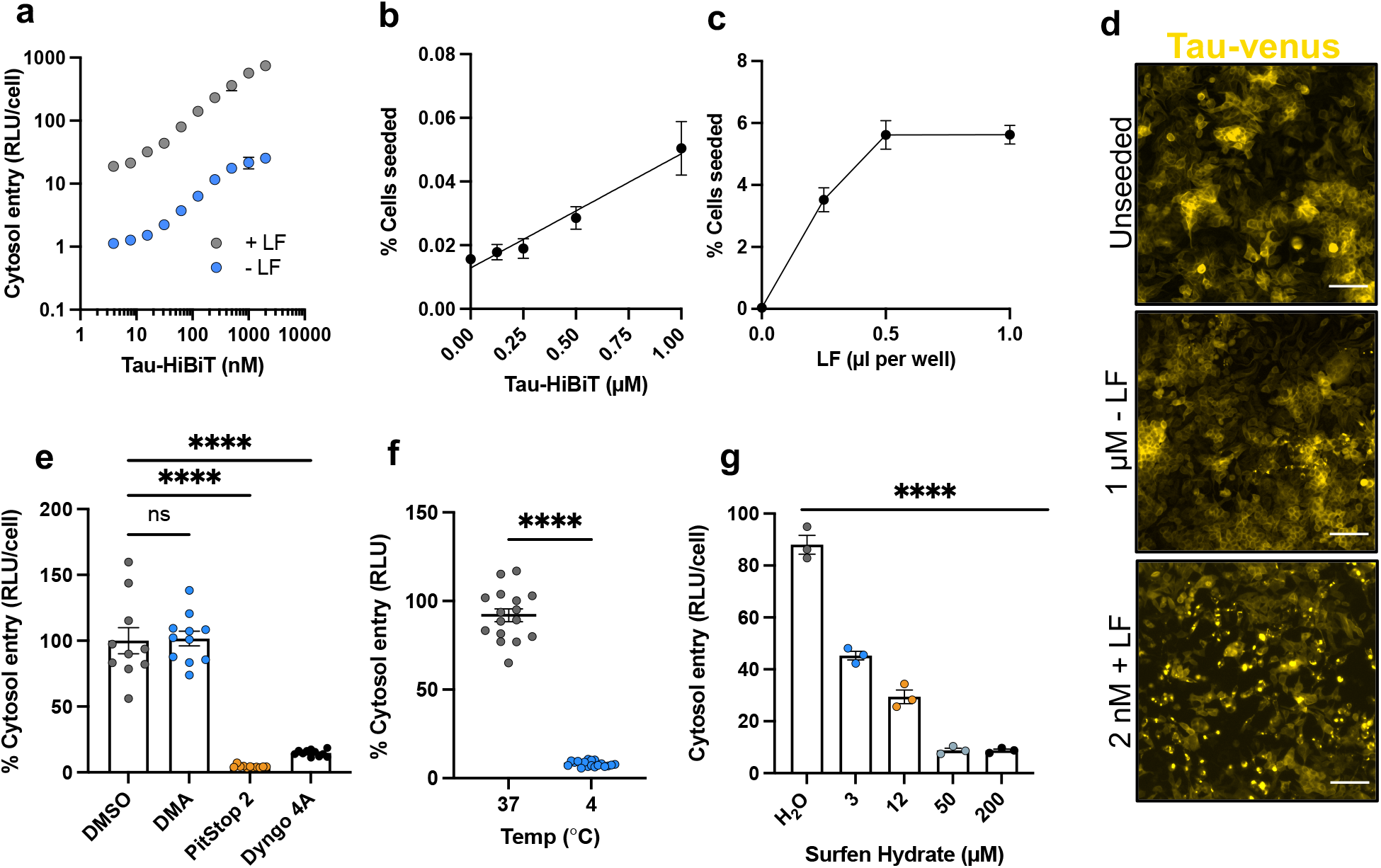
Tau entry relies on clathrin and dynamin dependent endocytosis. **a)** Entry curve of increasing concentrations of tau-HiBiT assemblies in the presence or absence of 1 µl LF after 24 h in NGL cells; n=3. **b)** Representative titration curve showing the effect of tau-HiBiT assembly concentration on seeded aggregation after 72 h in HEK293 cells expressing P301S tau-venus; n=3, 5 fields analysed per well. **c)** Representative curve showing the effect of increasing concentrations of LF on the seeded aggregation of 2 nM tau-HiBiT assemblies; n=3, 5 fields analysed per well. **d)** Fluorescence microscopy images of unseeded control (PBS treated) P301S tau-venus cells, cells seeded with 1 µM tau-HiBiT without LF, or cells seeded with 2 nM Tau-HiBiT in the presence of 0.5 µl LF. Scale bars, 100 µm. **e)** HiBiT assemblies. HEK-NGL cells were pre-treated with DMA (200 µM), PitStop 2 (10 µM) or Dyngo 4a (20 µM) for 30 minutes before challenge; n=≥3, N=3 independent biological replicates. **f)** Effect of temperature on tau entry. 50 nM tau-HiBiT assemblies were supplied to HEK-NGL cells for 1 h at 37°C or 4°C; n=16. **g)** The effect increasing concentrations of surfen hydrate on tau entry to HEK-NGL cells at 1 h. Cells were pre-treated for 30 minutes with indicated concentrations, or equivalent concentration of solvent (water) before 50 nM tau-HiBiT addition; n=3. DMA; dimethyl-amiloride, LF; lipofectamine 2000, RLU; relative light units. Error bars are mean ± s.e.m.

### Compromised endolysosomal machinery increases tau entry

After observing the dependency of tau entry on clathrin coat protein and dynamin GTPase, we next investigated the intracellular compartment from which tau assemblies escape. Clathrin- and dynamin-dependent endocytosis results in the enclosure of cargo in a primary endocytic vesicle, which undergoes homotypic fusion to form an early sorting endosome that subsequently traffics throughout the endosomal network with the intraluminal pH simultaneously decreasing^46^. Rab GTPases are crucial effectors of this endosome maturation. To investigate the nature of the compartment from which tau escapes in HEK293 cells, we used siRNA against early endosomal RAB5A or late endosomal RAB7A and performed tau entry assays over the course of 4 h (**Fig. 3a,b**). Interestingly, we observed an increase in entry when RAB7A, but not RAB5A, was depleted, suggesting a sensitivity of entry to impaired late endosomal compartments.

**Figure 3:**
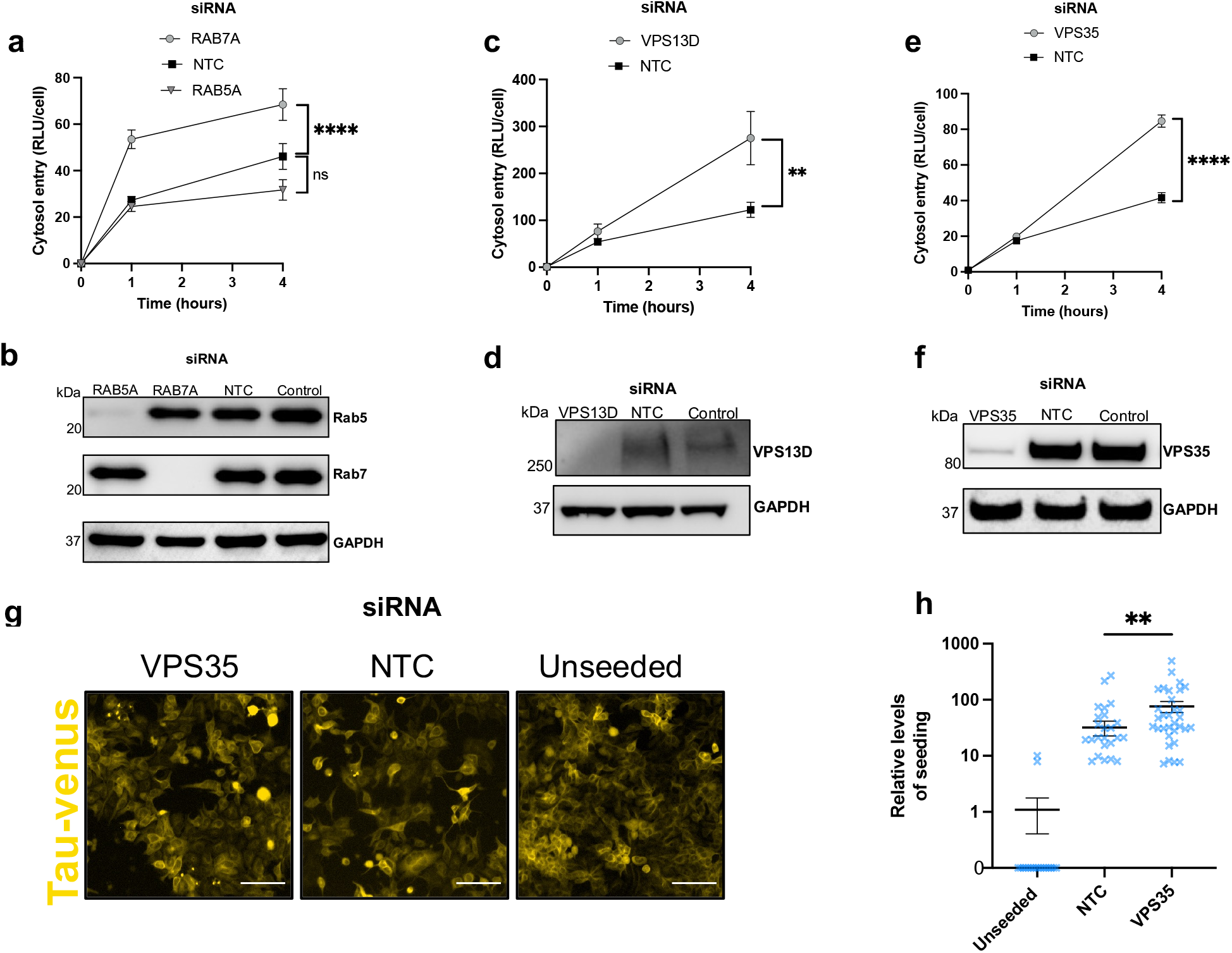
Endocytic machinery is required to prevent tau access to the cytosol. **a)** 4 h entry time course of 50 nM tau-HiBiT assemblies following RAB5A or RAB7A knockdown in HEK-NGL cells. Cells were treated with siRNA for 72 h before assaying tau entry; n=3, N=3 independent biological replicates. **b)** Western blot of cell lysates in (**a**) probing for Rab5, Rab7 and GAPDH **c)** 4 h entry time course of 50 nM tau-HiBiT assemblies in HEK-NGL cells following VPS13D knockdown. Cells were treated with siRNA for 72 h before assaying tau entry; n=3, N=3 independent biological replicates. **d)** Western blot of cell lysate from (**c**) probing for VPS13D and GAPDH. **e)** 4 h entry time course of 50 nM tau-HiBiT assemblies in VPS35 knockdown HEK-NGL cells. Cells were treated with siRNA for 72 h before assaying tau entry; n=3, N=2 independent biological replicates. **f)** Western blot of cell lysates in (**e**) venus cells treated with control (water treated) or VPS35-targeting siRNA 72 h after the addition of 250 nM exogenous tau-HiBiT assemblies in the absence of transfection reagents. Scale bars, 100 µm. **h)** Quantification of seeded aggregation from tau-venus seeding assays shown in panel **h**; n=6, 5 fields per well analysed. NTC; non-targeting control. RLU; relative light units. Control corresponds to untreated cells. Error bars are mean ± s.e.m.

Recently, it was shown that compromised function of VPS13D, a ubiquitin binding protein^47^, promoted the seeded aggregation of tau^48^. It was proposed that the potential mechanism of augmented tau aggregation may be a result of increased endolysosomal escape of tau seeds. To determine whether VPS13D is involved in maintaining tau assemblies from accessing the cytosol, we depleted VSP13D in HEK-NGL cells and observed a marked increase in tau entry (**Fig. 3c,d**). This corroborates the finding that VPS13D is essential to maintaining vesicular integrity and confirms that its depletion promotes entry. We next investigated the role of VPS35, a core constituent of the endosome-to-golgi retromer complex, an essential component of proper endosome sorting. Deficiencies in VPS35 are linked to increased tau burden and late-onset AD^49,50^ as well as being a known genetic risk in Parkinson’s disease^51^. We treated cells with VPS35 siRNA and, similar to VPS13D depletion, we observed a significant increase in tau entry in knockdown cells but not in control treated cells over 4 h (**Fig. 3e,f**). Given that VPS35 interacts with RAB7^53^, and that genetic silencing of VPS35 increases tau accumulation^49^, our results suggest a mechanism by which mutations associated with the gene may alter the progression of tau pathology. To further support our observations in the entry assay, we depleted VPS35 in HEK293 cells expressing P301S tau-venus and performed seeding assays in the absence of transfection reagents. We observed a significant increase in fluorescent puncta corresponding to tau aggregation in VPS35-depleted cells relative to control cells (**Fig. 3g,h**). These data demonstrate an essential role for endosome sorting and repair machinery in preventing tau escape to the cytosol.

### The mechanism of tau entry is cell-type dependent

To investigate the mechanism of tau entry in a more physiologically relevant system, we adapted our assay to primary neurons derived from wild-type (WT) C57BL/6 mice. We used chimeric particles of adeno-associated virus produced with capsids of types 1 and 2 (AAV1/2) to deliver a self-cleaving variant of the LgBiT reporter from a neuron-specific synapsin (hSyn) promoter relying on the T2A peptide^54^. This construct, hSyn::eGFP-P2A-LgBiT-NLS (GPLN), provides eGFP as a fluorescent marker and low levels of nuclear-localised LgBiT (**Fig. 4a-d**). An optimised protocol was established with AAV challenge at 2 days in vitro (DIV) for tau entry assays at 7 DIV, where we see close to 100% of neurons expressing GPLN (**Fig. 4d, Fig. S2a-c**).

**Figure 4:**
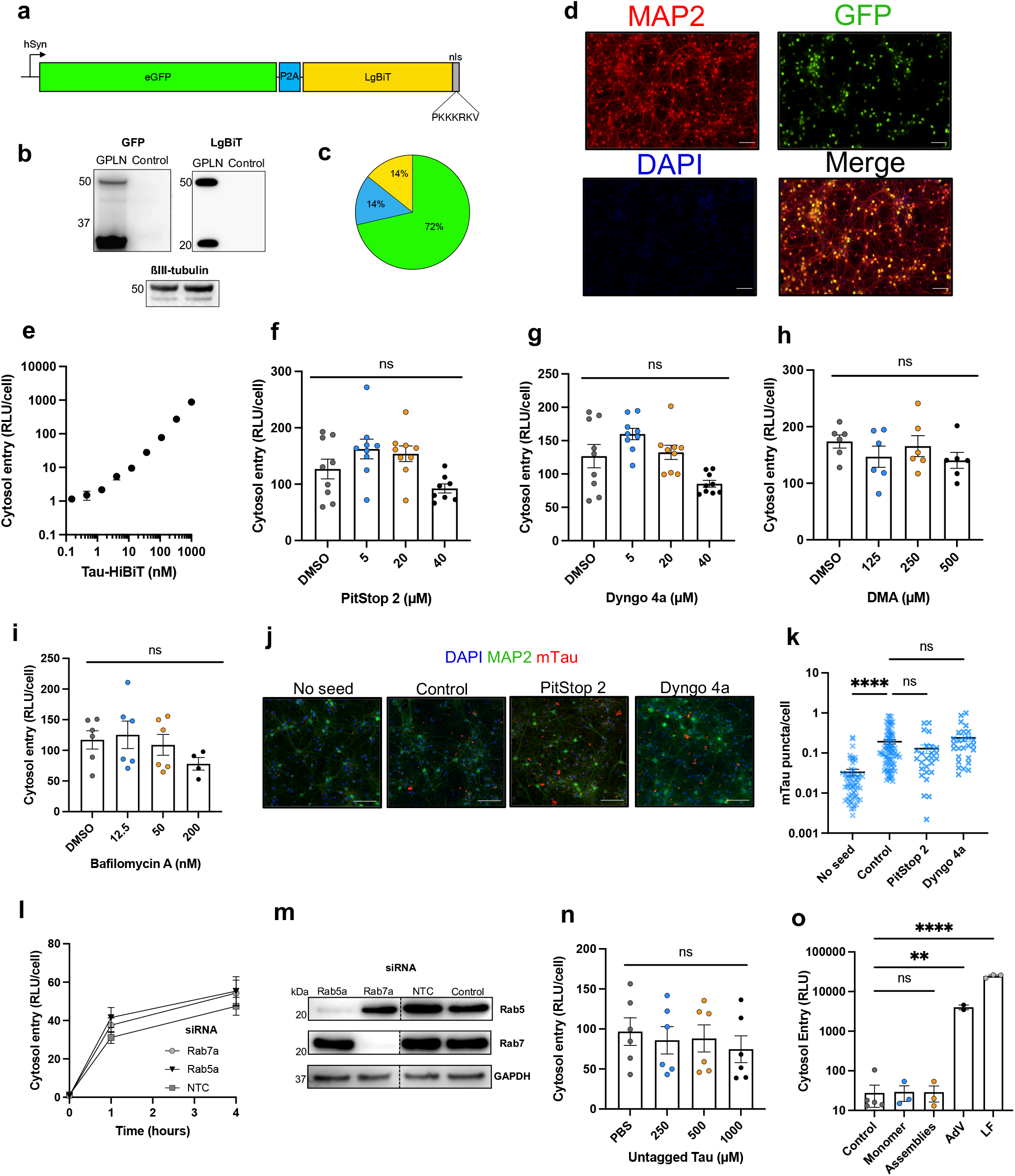
Tau entry to mouse primary neurons is clathrin and dynamin independent and is not mediated by tau itself. **a**) Cartoon depicting the GPLN construct, to scale relative to GFP. **b**) Western blotting 7 DIV WT neurons 5 days post-challenge with AAV1/2-hSyn-GPLN, probing for GFP, LgBiT, and βIII-tubulin as a neuronal loading control. **c**) Pie chart visualising the proportion of protein products translated from the GPLN construct. Green, GFP only; blue, GFP-P2A-LgBiT-nls; yellow, LgBiT-nls only. **d**) Fluorescence microscope images of WT 7 DIV primary neurons transduced with AAV1/2-hSyn-GPLN. Scale bars, 100 µm. **e**) Representative 1 h entry titration curve after addition of tau-HiBiT assemblies onto WT 7 DIV primary neurons expressing GPLN; n=3. **f-i**) Effect of clathrin inhibition (PitStop 2), dynamin inhibition (Dyngo 4a) Na^+^\H^+^ import inhibitor DMA or vacuolar ATPase inhibitor bafilomycin A on neuronal tau entry. WT 7 DIV GPLN expressing neurons were pre-treated for 1 h with indicated compound and entry of 50 nM tau-HiBiT assemblies quantified after a further 1 h. n=≥3 from N=3 independent biological replicates (panel **f, g**) n=3 from N=2 independent biological replicates (panel **h, i**). **j**) Seeded aggregation of tau in DIV 14 WT neurons following pre-treatment for 1 h with control solvent (0.1% DMSO), PitStop 2 (10 µM), Dyngo 4a (2 µM) before challenge with 100 nM recombinant tau-HiBiT assemblies at 7 DIV. Drugs were left on cells for the 7-day period. Scale bars, 100 µm. **K**) Quantification of seeded aggregation shown in panel **j.** n=3, 15 fields per well analysed. **l**) 4 h entry time course of 50 nM tau-HiBiT assemblies in WT 7 DIV neurons expressing GPLN 72 h post-knockdown with 1 µM Rab5a, Rab7a or NTC siRNA; n=3, N=3 independent biological replicates**. m**) Western blotting of neuronal knockdown lysates from panel **L** for Rab5, Rab7, and GAPDH as a loading control. Dashed line represents membrane cropping, full blot available in supplementary information (**Fig. S2h**). (**n**) Effect of untagged tau on cytosolic entry of 50 nM tau-HiBiT assemblies to neurons. Untagged tau aggregates were co-applied to WT 7 DIV neurons expressing GPLN and entry quantified after 1 h; n=3, N=2 independent biological experiments. **o**) Endosomal lysis assay by delivery of Nluc plasmid to HEK293 cells in the presence of excess tau monomer or aggregate (1 µM), adenovirus (MOI 100 IU/cell) or LF (0.2 µl/well), n=3. AdV, adenovirus type 5; DMA, dimethyl amiloride; GPLN, eGFP-P2A-LgBiT-nls; NTC, non-target control; hSyn, human synapsin promoter; RLU, relative light units. Error bars are mean ± s.e.m.

We titrated tau assemblies onto GPLN-neurons and observed a concentration-dependent luciferase signal that was resistant to trypsin, confirming that luminescence originates from an intracellular tau-HiBiT:LgBiT interaction (**Fig. 4e, Fig. S2d**). Investigating the entry mechanisms of tau in neurons, we were surprised to find that neither clathrin nor dynamin inhibition with PitStop 2 or Dyngo 4a reduced the entry of tau to the cytosol (**Fig. 4f,g**). Similarly, treatment with DMA to block Na^+^/H^+^ import channels was unable to prevent entry (**Fig. 4h**). We also investigated the requirement for low endosomal pH in entry. We treated neurons acutely with the V-ATPase inhibitor bafilomycin-A at commonly used concentrations to inhibit acidification but avoid downstream effects on autophagy^55^. We found no requirement for V-ATPase mediated endosomal acidification in tau entry (**Fig. 4i)**. In all cases, neuronal viability was monitored to ensure that experiments reflect entry to healthy cells (**Fig. S2e**). We confirmed that seeded aggregation is similarly insensitive to these inhibitors by challenging DIV 7 neurons with 100 nM tau-HiBiT assemblies in the presence or absence of PitStop 2 or Dyngo 4a for 7 days. We observed formation of endogenous mouse tau (mTau) aggregates in the presence of inhibitors which showed no significant deviation from the solvent treated control neurons (**Fig. 4j,k**). Given that tau can spread in a transsynaptic manner^56–58^, we considered whether the observed independence was a result of performing entry assays in cells with improperly formed synapses, which occurs around 7-10 DIV^59,60^. We therefore performed entry assays in DIV 14 neurons but consistently found no role for clathrin or dynamin in entry of tau to the cytosol of neurons (**Fig. S2f**). To further interrogate the entry pathway of tau to neurons, we knocked down mouse Rab5a or Rab7a via siRNA and monitored tau-HiBiT entry over 4 h (**Fig. 4l**). Despite a knockdown efficiency of >90% for both proteins and minimal toxicity, we found no role for either protein in tau entry to neurons (**Fig. 4m, Fig. S3g,h**). Our results suggest that tau entry to the cytosol of neurons occurs via an alternative mechanism, independent of clathrin-mediated endocytosis. The results in neurons are in direct contrast to observations in HEK293 cells, suggesting profound differences in the underlying biology between these systems.

### Tau assemblies do not mediate their own escape to the cytosol

One possibility is that tau assemblies mediate their own escape from vesicles following uptake by destabilising endosomal membranes. To test this, we titrated tagless tau assemblies in the presence of a constant (50 nM) concentration of tau-HiBiT. If the tagless tau assemblies promoted membrane rupturing, we predicted this would result in an increase in tau-HiBiT entry. However, in both HEK293 cells and primary neurons, we found no such increase in signal, suggesting that tau does not mediate its own entry (**Fig. 4n, Fig. S3a**). We then used a second membrane integrity assay wherein a plasmid encoding luciferase is supplied to the extracellular media of HEK293 cells. Addition of membrane rupturing agents results in plasmid transfer to the intracellular environment, promoting luciferase expression. The addition of LF or human adenovirus type 5, which enters cells via endosomal rupture, resulted in 100- to 1,000-fold increases in the levels of luminescence. In contrast, tau assemblies and tau monomer at high concentration (1 µM) resulted in no observable signal above background (**Fig. 4o**).

### Lrp1 and HSPGs facilitate the cholesterol-dependent entry of tau to the cytosol of neurons

We next investigated the receptor-dependency of tau entry. We first depleted the low density lipoprotein receptor, Lrp1, recently identified as a receptor for tau uptake^13^. Knockdown of this protein substantially decreased tau entry to cells at 4 h after addition (**Fig. 5a,b**). However, entry at 1 h was less affected by Lrp1 depletion, potentially reflecting residual membrane Lrp1 remaining available for uptake followed by receptor internalisation upon tau binding. Addition of heparin to the cell media reduced tau entry in a dose-dependent manner, consistent with a role for cell-surface HSPGs in promoting tau attachment and uptake to neurons (**Fig. 5c**). As cholesterol biology is heavily implicated in susceptibility to tauopathies, we next investigated the dependency of cholesterol in neuronal tau entry.

**Figure 5:**
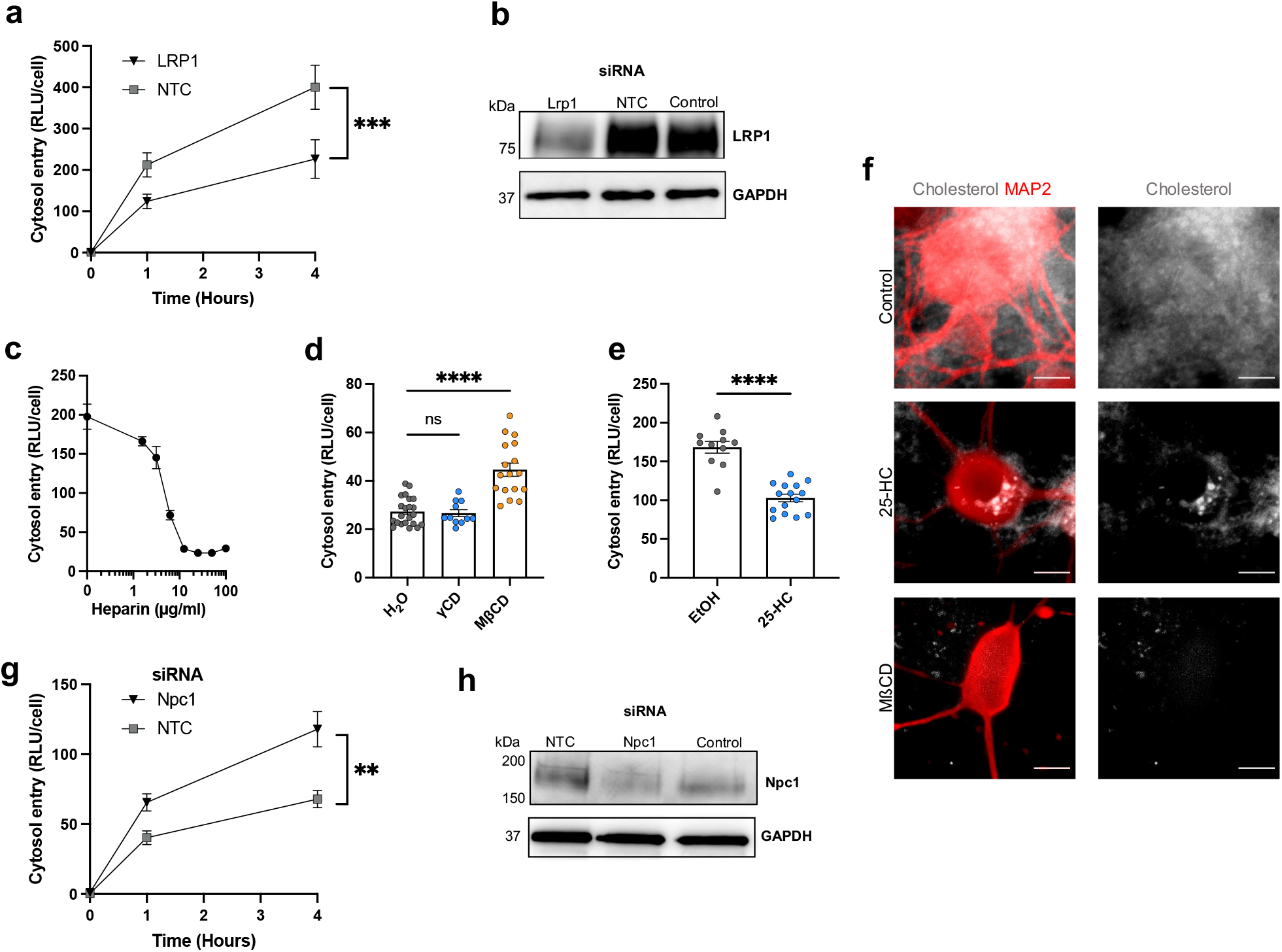
Tau entry to mouse primary neurons depends on receptor-mediated uptake and is controlled by cholesterol. **a)** Time-course of entry of 50 nM tau-HiBiT assemblies in WT DIV 7 neurons expressing GPLN 72 h post-knockdown with Lrp1 or NTC siRNA; n=3, N=3 independent biological replicates. **b)** Western blotting of neuronal lysates from panel **a** for Lrp1 and Gapdh as a loading control. **c)** Representative entry curve of 50 nM tau-HiBiT assemblies in 1 h to neurons in the presence of increasing concentrations of heparin; n=3. **d**) Effect of cholesterol extraction on entry of 50 nM tau-HiBiT assemblies in WT DIV 7 neurons expressing GPLN after 1 h. Neurons were pre-treated with γCD (2 mM) or MβCD (2 mM) for 2 h before addition of tau-HiBiT assemblies for 1 h; n=≥4, N=3 independent biological replicates. **e)** Effect of cholesterol transport inhibition on the entry of 50 nM tau-HiBiT assemblies over 1 h in WT 7 DIV neuron expressing GPLN. Cells were pre-treated for 16 h with 25-HC (10 µM) before the addition of tau-HiBiT assemblies for 1 h; n=≥3, N=3 independent biological experiments. **f)** Filipin III staining of WT 7 DIV neurons to visualise cellular cholesterol in control (0.1% EtOH), MβCD, (2mM, 2 h) or 25-HC (10 µM, 16 h) cells. Scale bar, 10µm. **g)** Entry time course of 50 nM tau-HiBiT assemblies in WT DIV 7 neurons expressing GPLN 72 h post-knockdown with Npc1 or NTC siRNA; n=3, N=3 independent biological replicates. **h**) Western blotting of neuronal lysates 72 h post knockdown with 1 µM NTC or Npc1 siRNA. Control corresponds to untreated cells. 25-HC; 25-hydroxycholesterol, MβCD; methyl-beta-cyclodextrin, GPLN; eGFP-P2A-LgBiT-nls, Npc1; Niemann-Pick C1 protein, NTC; non-targeting control, RLU; relative light units. Error bars are mean ± s.e.m.

We hypothesised that changes in membrane cholesterol may modulate the entry of tau to the cytosol. First, we treated neurons with the cholesterol extracting agent methyl-beta-cyclodextrin^61^ (MβCD). Doses in excess of 3 mM of this compound were found to be cytotoxic to neurons (**Fig. S4a**). However, treatment with 2 mM MβCD for 2 h before addition of 50 nM tau-HiBiT for 1 h was not toxic but effectively extracted cholesterol from neurons, as visualised by staining with the cholesterol-binding dye, filipin (**Fig. 5f, Fig. S4b**). These conditions resulted in a significant increase in tau entry to neurons (**Fig. 5d**). We verified that the effect was due to cholesterol extraction and not a result of off-target effects of cyclodextrins, as treatment with gamma-cyclodextrin (γCD), which does not extract cholesterol from membranes, resulted in no significant change in entry compared to the control.

Treatment of cells with the compound 25-hydroxycholesterol (25-HC) results in the accumulation of cholesterol in intracellular membrane-bound organelles^62^. 25-HC has been shown to inhibit entry of viruses to the cell, including SARS-CoV-2^63,64^. We pre-treated neurons for 16 h with 25-HC. Filipin staining revealed accumulation of intracellular cholesterol and we observed a significant reduction in entry (**Fig. 5e,f**). Mutations in the Niemann-Pick C1 protein (NPC1) are causative of NPC^65^, a primary tauopathy with mis-sorted cholesterol. We questioned whether depletion of Npc1 in primary neurons could modulate tau pathology, potentially informing the mechanism of action of NPC1 mutation in human disease. Npc1 depletion by siRNA resulted in a significant increase in tau entry (**Fig. 5g,h**). These results suggest that intracellular cholesterol transport and membrane cholesterol content are critical to the entry pathway of tau to the cytosol, and may therefore potentially modulate seeded aggregation of tau.

### Seeded aggregation of tau is sensitive to membrane cholesterol

Our findings have revealed the relationship between cellular co-factors and tau entry to the cytosol in immortalised cells and primary culture models. We next sought to investigate whether changes in tau entry leads to changes in seeded aggregation in a physiological setting. For these experiments, we used organotypic hippocampal slice cultures (OHSCs) which maintain authentic neural architecture and cell-type diversity^66^. We prepared OHSCs from P301S tau transgenic mice, which develop intracellular tau inclusions by 6 months of age^67^. OHSCs prepared from these mice do not develop detectable tau pathology unless challenged with tau assemblies^68^. Tau assemblies were provided to OHSCs following a 1-day pre-treatment period with endocytosis inhibitors or cholesterol modulators. Tau assemblies and drug were removed after 3 days and the OHSCs were incubated for a further 3 weeks before detection and quantification of seeded aggregation using the phospho-tau specific antibody AT8 (**Fig 6a-f**). Consistent with our observations for tau entry to the cytosol, inhibition of clathrin or dynamin with PitStop 2 or Dyngo 4a resulted in no significant change in seeded aggregation (**Fig. 6a**). Supporting our proposed role for cholesterol in the maintenance of the unseeded state, extraction of cholesterol with MβCD substantially increased seeded aggregation whereas the γCD control yielded no statistically significant change (**Fig. 6b**). Moreover, treatment with 25-HC resulted in a significant reduction in seeded aggregation, again consistent with its effect on tau entry (**Fig. 6c**).

**Figure 6:**
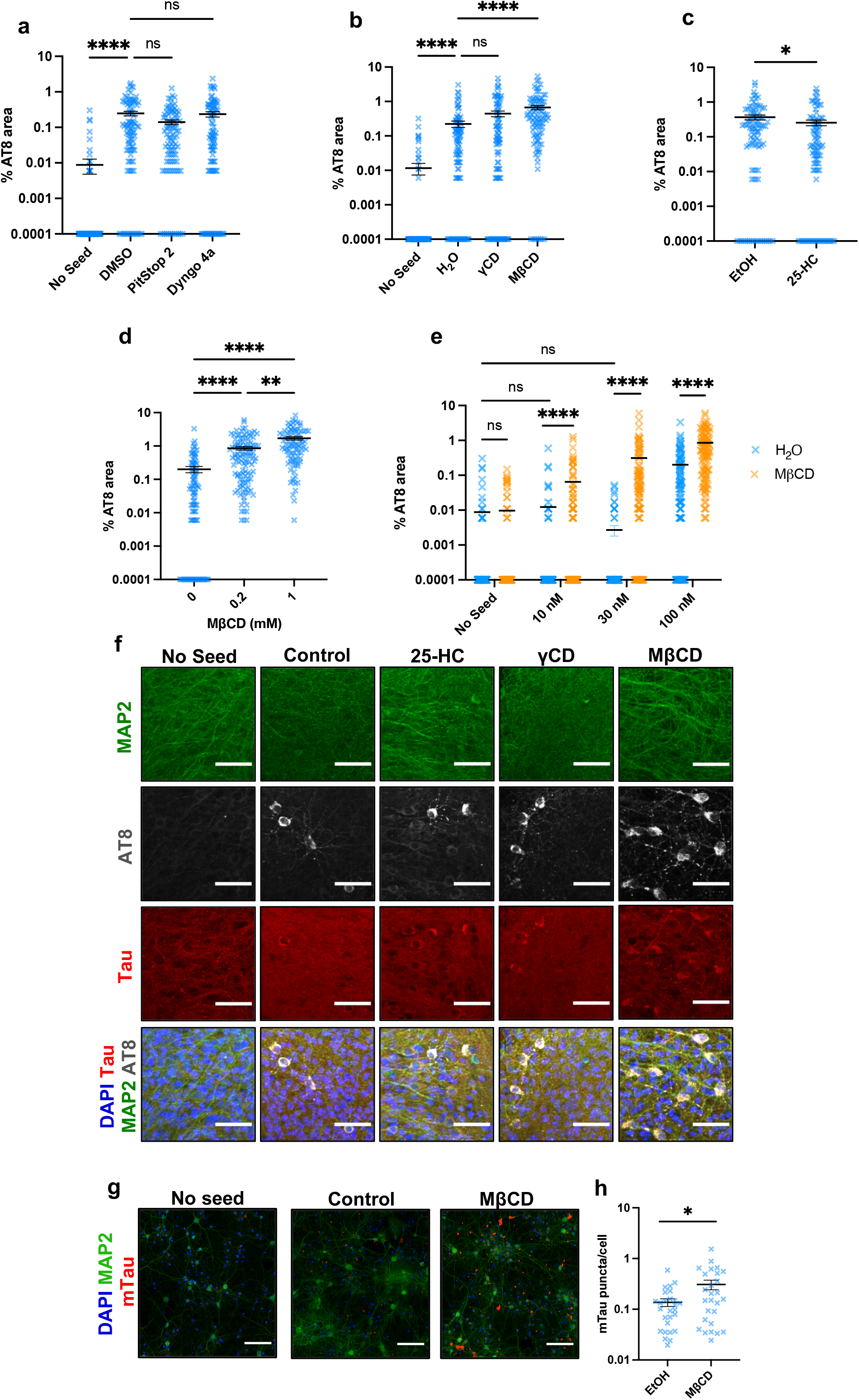
Cholesterol extraction promotes seeded aggregation in organotypic hippocampal slice cultures. **a)** Quantification of seeded aggregation in OHSCs from P301S tau transgenic mice pre-treated with control solvent (0.1% DMSO), Pitstop 2 (10 µM) or Dyngo 4a (2 µM) for 24 h prior to the addition of fresh drug and 100 nM tau assemblies for 3 days. Media was exchanged and OHSCs were incubated for a further 3 weeks prior to analysis by immunofluorescence microscopy; slices from N=6 mice per condition. **b)** Quantification of seeded aggregation in OHSCs from P301S tau transgenic mice pre-treated with control solvent (H20), γCD (200 µM) or MβCD (200 µM) for 24 h prior to the addition of fresh drug and 100 nM tau assemblies; slices from N=6 mice per condition. **c)** Quantification of seeded aggregation in OHSCs from P301S tau transgenic mice pre-treated with control solvent (0.1% EtOH) or 25-HC (10 µM) for 24 h prior to the addition of fresh drug and 100 nM tau assemblies; slices from N=6 mice per condition. **d)** Quantification of seeded aggregation in OHSCs from P301S transgenic mice treated with 100 nM tau assemblies in the presence of 200 µM or 1 mM MβCD. Slices from N=6 mice per condition. **e)** Titration curve of tau in the presence of MβCD (200 µM) or H20 as a solvent control. Seeded aggregation was quantified in OHSCs from P301S tau transgenic mice. Slices from N=6 mice per condition. **f)** Representative images of OHSCs quantified in panel **a-c**. Scale bars, 50 µm**. g**) Seeded aggregation of tau in DIV 14 WT neurons following pre-treatment with control solvent (0.1% EtOH) or MβCD (200 µM, 2 h) and challenge with 100 nM recombinant tau-HiBiT assemblies at 7 DIV. Drugs were left on cells throughout the 7-day period. n=3, Scale bars, 100 µm. **h)** Quantification of seeded aggregation shown in panel **g**. n=3, 15 fields per well analysed. Statistical analysis was performed via the Kruskal-Wallis test by ranks, and Dunn’s post hoc multiple comparisons test except for panel **e** which was analysed via multiple nonparametric Mann-Whitney tests. γCD; gamma-cyclodextrin, MβCD; methyl-beta-cyclodextrin, 25-HC; 25-hydroxycholesterol. Error bars are mean ± s.e.m.

We have previously shown that seeded aggregation in mouse neural tissue conforms to non-linear dynamics, with low levels of extracellular tau assemblies failing to produce detectable seeded aggregation^68^. We titrated tau assemblies in the presence or absence of 200 µM MβCD in OHSCs and observed a significant increase in tau entry when MβCD was present (**Fig. 6d, Fig. S5a**). Remarkably, seeded aggregation was observed at 10 and 30 nM supplied tau when MβCD was present. These concentrations completely failed to induce detectable seeded aggregation in slices that had not been treated with MβCD. We also found that aggregation-promoting effect of MβCD is titratable, as 1 mM MβCD further increased seeded aggregation relative to lower concentrations (**Fig. 6e**). These data taken together demonstrate an essential role of cellular cholesterol levels in maintaining membrane integrity in mouse neurons and neural tissue. Moreover, they suggest that dysregulated cholesterol permits seeded aggregation of tau when neurons are exposed to levels of extracellular tau that would otherwise remain safely in the extracellular environment.

We next tested whether aggregation of endogenous mouse tau was sensitive to cholesterol levels, by performing seeding assays in primary non-transgenic C57BL/6 mouse neurons. Cholesterol was extracted with a low, tolerated, concentration of MβCD, which potentiated seeding of endogenous mouse tau (**Fig. 6g, h)**. These findings demonstrate that seeding of both transgenic human tau and endogenous tau are sensitive to cholesterol levels in mouse primary neurons.

### Tau enters human neurons in a cholesterol sensitive manner

We next asked whether the tau entry pathway to mouse neurons is conserved with humans. We generated human cortical neurons derived from pluripotent stem cells (iNeurons) using a robust protocol^69^. Cytosolic LgBiT was expressed by transduction with AAV1/2-hSyn-GLPN. We verified LgBiT expression by western blot and flow cytometry, where we observed >95% transduction efficiency (**Fig. 7a, S6a**). We confirmed that day 14 iNeurons expressed essential neuronal and synaptic genes indicative of fully differentiated human cortical neurons (**Fig. 7b, S6b**). We first investigated the entry dynamics of tau assemblies in day 14 LgBiT expressing iNeurons (GPLN-iNeurons), where entry occurred in a concentration- and time-dependent manner, similar to the other cellular systems we have studied (**Fig. 7c, d).** We next questioned whether the human neurons would recapitulate the clathrin and dynamin dependence observed in human immortalised cell lines, or conversely display the endocytosis independence observed in mouse neurons. We pre-treated GPLN-iNeurons with clathrin inhibitor PitStop 2 or dynamin inhibitor Dyngo 4a and performed a 1 h entry assay with 50 nM tau-HiBiT assemblies. We found no role for clathrin or dynamin in tau entry in human neurons, consistent with observations in mouse neurons and OHSCs (**Fig. 7e)**. Similar to both mouse neurons and human immortalised cell lines, we found a strong dependence on HSPGs, as pre-treatment with 20 µg/ml heparin blocked entry by an average of ∼67% (**Fig. 7f**). We then investigated the cholesterol sensitivity of tau entry in human neurons, and we found that cholesterol extraction by pre-treating for 2 h with MβCD results in a >2-fold increase in entry compared to untreated cells. Again, we observed no significant change in entry with the inactive cyclodextrin, γCD (**Fig. 7g**). Finally, we pre-treated GPLN-iNeurons with 25-hydroxycholesterol and we saw a significant reduction in entry of an average of ∼47% (**Fig. 7h**). These data further support our findings from mouse model systems and corroborate the role of cholesterol in tau entry and seeded tau aggregation.

**Figure 7:**
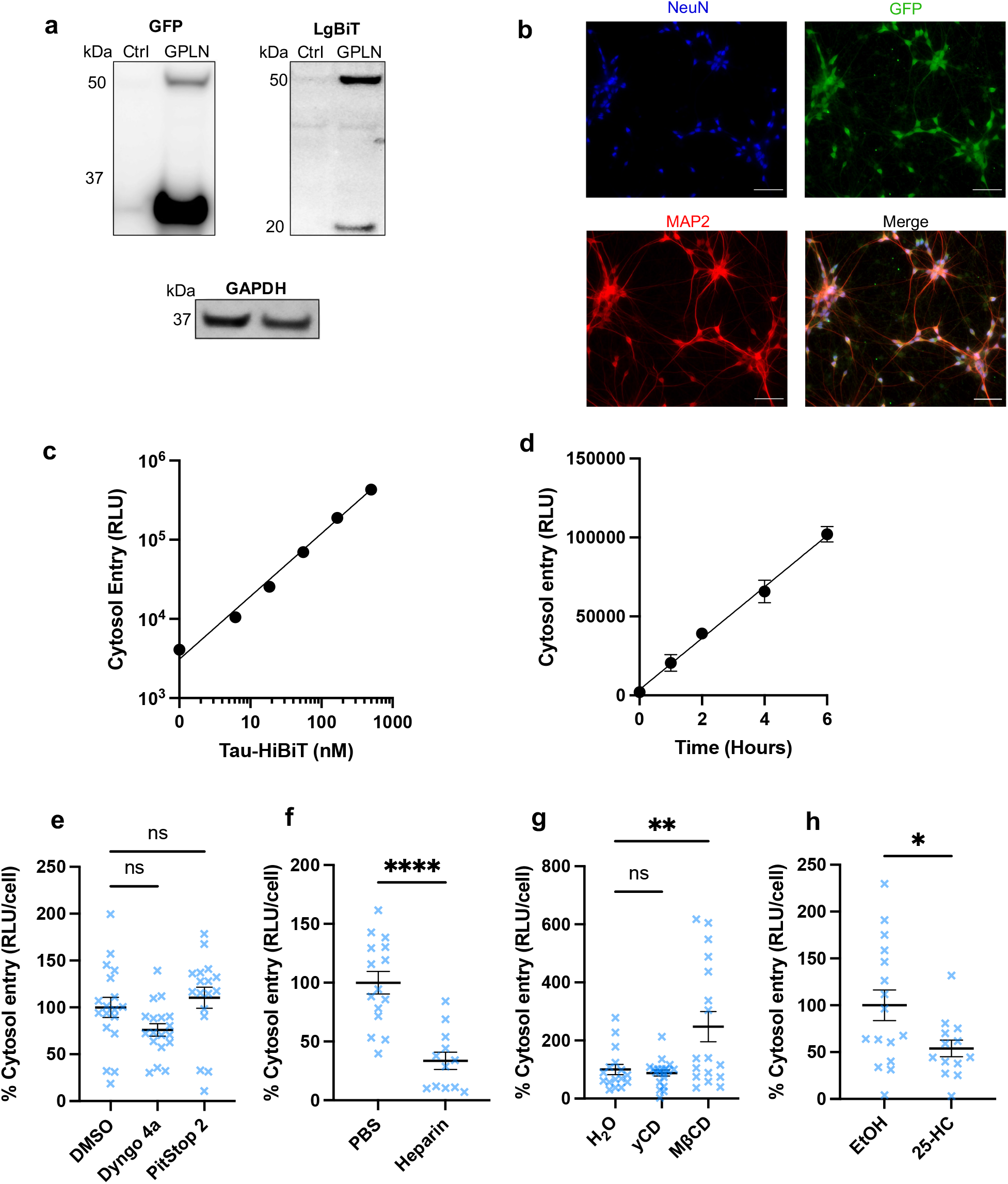
Tau enters human neurons in a cholesterol-sensitive manner. **a)** Western blotting of day 14 human neurons transduced with AAV1/2-hSyn-GFP-P2A-LgBiT-nls (GPLN-iNeurons) 7 days post challenge probing for LgBiT, GFP and GAPDH as a loading control. **b)** Representative fluorescence microscopy images of DIV 14 GPLN-iNeurons stained for MAP2, GFP, and the neuronal nuclear antigen NeuN. Scale bars, 50 µm. **c)** Representative titration curve of tau-HiBiT assemblies added to DIV 14 GPLN-iNeurons and entry assayed after 1 h, n=3. **d)** Representative time-dependency curve when 50 nM tau-HiBiT assemblies were added to GPLN-iNeurons and entry assayed after 1, 2, 4 and 6 h, n=3. **E)** Effect of clathrin and dynamin inhibition on tau entry in GPLN-iNeurons. Neurons were pre-treated with PitStop 2 (30 µM), Dyngo 4a (30 µM) or DMSO (0.1%) for 1 h prior to assaying the entry of 50 nM tau-HiBiT assemblies in 1 h. n=3-6 from N=3 independent differentiations. **f)** Effect of HSPG inhibition on tau entry in GPLN-iNeurons. Neurons were pre-treated with heparin (1 µM) or PBS for 1 h prior to assaying the entry of 50 nM tau-HiBiT assemblies in 1 h, n=3-6 from N=3 independent differentiations. **g)** Effect of cholesterol extraction on tau entry in GPLN-iNeurons. Neurons were pre-treated with cholesterol extracting reagent MβCD, control cyclodextrin γCD or H2O for 2 h prior to assaying the entry of 50 nM tau-HiBiT assemblies to the cytosol in 1 h, n=6 from N=3 independent biological experiments. **h)** Effect of cholesterol transport inhibition on tau entry in GPLN-iNeurons. Neurons were pre-treated with 25-hydroxycholesterol (10 µM) or EtOH (0.1%) for 16 h prior to assaying the entry of 50 nM tau-HiBiT assemblies in 1 h. n=4 from N=3 independent differentiations. GPLN; eGFP-P2A-LgBiT-nls, MβCD; methyl-beta-cyclodextrin, 25-HC; 25-hydroxycholesterol, RLU; relative light units. Error bars are mean ± s.e.m.

## Discussion

In this study we have developed sensitive assays capable of detecting the entry of tau to the cytosol following its application at nanomolar concentrations to the cell exterior. The high sensitivity of the assay means that entry biology can be studied without the use of extra-physiological concentrations of tau. Using these assays, we have described the entry of tau to the cytosol in two cell-based models: HEK293 cells, which are widely used as reporters for seeded aggregation, and neurons, the major site of tau pathology in the brain of AD patients. Entry occurred in a dose-dependent manner over several orders of magnitude, suggesting that extracellular tau concentrations contribute linearly to entry. Levels of entry in HEK293 and neurons were a critical determinant for seeded aggregation, consistent with entry, rather than uptake, being the rate-limiting step to seeded aggregation. We thus quantitatively described a link between entry and seeded aggregation and establish a causal relationship between these processes.

### Cytosolic entry of tau is cell-type dependent

Surprisingly, we observed that the mechanism of entry was highly divergent between HEK293 cells and neurons. We found that tau entry in HEK293 cells is dependent on the coat protein clathrin and the GTPase dynamin, consistent with the frequently described role of canonical endocytosis in human immortalised cell lines uptake^17^. When we induced endolysosomal dysfunction in HEK293 cells via genetic knockdown of critical endosomal genes RAB5A and RAB7A, we found only knockdown of RAB7A increased entry. Furthermore, knockdown of VPS35, a core constituent of the retromer complex and a RAB7 effector, similarly increased entry. We therefore consider it likely that tau escapes from late endosomal compartments in HEK293 cells. This dependency was lost, however, in primary mouse neurons. Here, entry of tau occurred through a clathrin- and dynamin-independent pathway with no observable effect of Rab5 or Rab7 depletion. This was not an artefact of using neurons with immature synaptic biology, as 14 DIV cells with highly connected morphology displayed a similar phenotype. Clathrin- and dynamin-independent endocytosis has been described for particular substrates suggesting that tau entry occurs from a specific subset of endosomal compartments ^70^, or potentially at the plasma membrane. Further supporting these findings, cytosolic entry of tau also occurred in a clathrin- and dynamin-independent manner in iPSC derived human neurons which form highly connected synaptic networks and express an array of mature neuronal markers. This therefore indicates that this mechanism is a result of neuron biology, rather than mechanism specific to mouse systems. Previous studies that have examined the uptake of tau to neurons have demonstrated involvement of clathrin-mediated endocytosis^14^, macropinocytosis^71^ and a clathrin-independent, dynamin-dependent pathway^72^. It is therefore possible that several pathways are responsible for tau uptake but tau escape to the cytosol occurs from a subset of compartments. This raises the prospect that molecular dissection of the entry pathway may permit the identification of inhibitors that prevent seeded aggregation by blocking entry of tau to the cytosol.

### Cholesterol is a critical determinant of tau entry

Cholesterol homeostasis has been repeatedly implicated in the risk and pathogenesis of Alzheimer’s disease^73^. Polymorphisms in APOE, the transporter for low density lipoprotein, are the main genetic risk factors for Alzheimer’s disease, with the E4 variant increasing the genetic risk in a dose-dependent manner. The involvement of APOE in AD has focussed predominantly on its effects on β-amyloid (Aβ). However, recent data demonstrate that APOE polymorphisms also directly impact tau pathogenesis. APOE polymorphisms affect the risk of progressive supranuclear palsy, which does not feature Aβ pathology^74^. Interestingly, homozygosity of the E2 variant, rather than E4, is the genotype associated with highest risk of tauopathy. Consistent with these studies, mouse models have revealed that APOE can modulate tau pathology independent of Aβ^75^. In the present study, we found that cholesterol levels in neurons are a critical determinant of tau entry. Cholesterol content alters the physiochemical properties of lipid bilayers, by increasing curvature rigidity and reducing the density of lipid packing^76^. Depletion of cholesterol using MβCD alters the mechanical parameters of membranes and renders them susceptible to rupture^77^. In contrast, 25-HC treatment increases the availability of cholesterol in phospholipid membranes^78^. In our experiments, extraction of cholesterol using MβCD resulted in increased entry and 25-HC reduced entry, consistent with adequate cholesterol content being essential for preventing tau entry. Of potential significance therefore, is the finding that cholesterol content of the plasma membrane is depleted in an age-dependent manner^23^. Given that age is the most important risk factor for dementia, the results potentially indicate a mechanism that permits tau pathology to accumulate in aged cohorts. Furthermore, identification of low cholesterol as a feature that renders neurons highly susceptible to seeded aggregation may permit future therapeutic intervention in tauopathies.

### Npc1 depletion enhances tau entry in neurons

95% of NPC patients harbour a loss-of-function mutations in NPC1, and many patients present with accumulation of tau filaments at an early age. In the present study, we found that depletion of the mouse homologue, Npc1, in primary neurons resulted in enhanced tau entry. Loss-of-function NPC1 mutations impair the lysosomal export of cholesterol and subsequent intracellular trafficking, and mutations in the sterol sensing domain of NPC1 result in impaired delivery of cholesterol to the plasma membrane^79^. Taken together with our findings that cholesterol extraction via MβCD results in enhanced entry and increased seeded aggregation, these data indicate that the cholesterol content of the neuronal plasma membrane is critical in maintaining the unseeded state. Further, the results provide a potential mechanistic underpinning to the observed accumulation of tau pathology in NPC patients.

### Tau escape to the cytosol

It has been suggested that tau escape to the cytosol is dependent on membrane destabilising activities of tau itself. This interpretation is based on the use of cytosolic galectin reporters, which bind to carbohydrates on the luminal leaflet of ruptured vesicles^15,56,80,81^. The addition of tau assemblies, or cell lysate containing tau assemblies, was found to increase the number of galectin-positive structures in those studies. Another study demonstrated that tau assemblies can bind to cell membranes via interactions between the repeat region and phospholipid bilayers, potentially consistent with direct lysis of membranes^82^. However, in our study, which directly measures entry to the cytosol, we found no evidence of tau-mediated entry in two separate assays and in both HEK293 and primary neurons. The reasons for these differences are not clear, though may reflect different cell types and fibril preparations, or may reflect inherent differences in directly measuring tau cytosolic entry versus galectin puncta. Our study used recombinant fibrils of a single isotype whereas fibrils prepared from the AD brain comprise all six CNS isoforms. We therefore cannot rule out that other isoforms or tau structures may bear membrane destabilising activity. However, the membrane-interacting domain of tau was mapped to the R2 repeat region^82^, which is present in the tau isoform used in the present study. Alternatively, membrane destabilisation may be a property of small oligomers, or monomers, rather than the filamentous assemblies used here^83,84^. Our results nonetheless favour an alternative model in which cellular cofactors, rather than tau, are the critical determinants in maintaining the integrity of membrane-bound compartments. Loss of this homeostatic control results in increased entry of tau to the cytosol and a concomitant increase in seeded aggregation. This suggests that cellular/genetic determinants will likely play a pivotal role in this rate-limiting step to tau pathology. The potential dynamic range of this rate-limiting step seeding is demonstrated by lipofectamine, which increased entry and seeded aggregation by >50-fold. In support of this interpretation, Chen et al. also found no evidence of tau mediating its own escape to the cytosol and instead suggested genetic factors influence membrane integrity^48^. Their results implicated vacuolar sorting proteins as playing a role in maintaining tau out of the cytosol, which we confirm here. Genome-wide association studies and certain dominantly inherited proteopathies implicate several endosome-associated proteins in neurodegeneration^85–87^. Early endolysosomal genes such as BIN1, PICALM and CD2AP are among the most common genetic polymorphisms associated with late-onset AD, yet there is no consensus on how they influence AD^88,89^. Future work should therefore ascertain whether membrane integrity is governed by such genetic risk factors and whether this predisposes to the transit of tau assemblies between cells of the brain.

## Summary

Our study provides the first quantitative description of tau entry to the cytosol. We establish that entry is essential for and rate-limiting to seeded aggregation of intracellular tau pools. Tau transits to the neuronal cytosol in a cholesterol sensitive manner following interactions with HSPGs and Lrp1. Entry is independent of Rab5/Rab7 GTPases and clathrin in neurons, suggesting that entry occurs from a specific subpopulation of vesicles that lack these cofactors. Dysregulated cholesterol, by extraction with MβCD or through Npc1 depletion, renders neurons vulnerable to tau entering the cytosol and permits seeded aggregation at concentrations of extracellular tau assemblies that fail to induce seeding in untreated cells. Our study enables the investigation of tau entry as distinct from uptake and will permit investigation of the genetic and pharmacologic factors that modify this rate-limiting step in seeded tau aggregation.

## Acknowledgements & Funding

We gratefully thank Dr Michel Goedert for provision of P301S tau transgenic mice and Prof Steven Paul for insightful conversations. We thank Dr Filomena Gallo of the Cambridge Advanced Imaging Centre for support and assistance in this work. We thank Annabel Smith and Sophie Sanford of the Cambridge UK DRI for preparation of tau assemblies. WAM is a Lister Institute Fellow and supported by a Sir Henry Dale Fellowship jointly funded by the Wellcome Trust and the Royal Society (Grant No. 206248/Z/17/Z). This work was supported by the UK Dementia Research Institute which receives its funding from DRI Ltd, funded by the UK Medical Research Council, Alzheimer’s Society and Alzheimer’s Research UK. BJT is supported by the Cambridge Trust Vice Chancellor’s Award and Hughes Hall Edwin Leong PhD scholarship. This project has received funding from the Innovative Medicines Initiative 2 Joint Undertaking under Grant agreement No. 116060 (IMPRiND). This Joint Undertaking receives support from the European Union’s Horizon 2020 research and innovation programme and EFPIA. This work is supported by the Swiss State Secretariat for Education, Research and Innovation (SERI) under contract number 17.00038. Schematic diagrams were prepared in BioRender.

## Author contributions

WAM and BJT conceived the study, designed experiments, and wrote the manuscript. BJT performed the majority of experiments with further experiments performed by LCVM, ELW, TK and WAM. SC, MJV, CK, LT, EM and LCJ provided reagents and prepared materials essential to the study. All authors edited the manuscript.

## Materials and Methods

### Antibodies

The following rabbit polyclonal antibodies were acquired from Proteintech; RAB5A (11947-1-AP), RAB7A (55469-1-AP), GFP tag (50430-2-AP), and tubulin (11224-1-AP) as well as secondary goat anti-mouse-HRP (SA00001-1) and goat anti-rabbit-HRP (SA00001-2). LgBiT antibody (N710A) and HiBiT antibody were acquired from Promega. Chicken polyclonal MAP2 (ab5392), rabbit monoclonal histone H3 (ab176842) and rabbit polyclonal VPS13D (ab202285) were purchased from Abcam. Mouse-monoclonal VPS35 (sc-374372) and cyclophilin-B (sc-130626) were purchased from Santa Cruz Biotech. Pan-tau (A0024) DAKO was acquired from Agilent. Anti-mouse tau (T49) was purchased from Sigma (MABN827). anti-phospho tau (AT8) (MN1020), Mouse monoclonal GAPDH loading control (MA5-15738) and Alexa-fluor conjugated secondary antibodies were purchased from Thermo Fisher.

### Plasmids

The following plasmids were purchased from addgene: pAAV-hSyn-eGFP (#50465), pAdDeltaF6 helper (#112867), pAAV 2/1 (#112862), pAAV 2/2 (#104963). pcRV-Gag-Pol was a gift from Prof Stuart Neil, Kings College London. pMD2G was a gift of Prof Didier Trono, EPFL. HIV-1 expression vector pPMPP was created by replacing the SFFV promoter from pSMPP with a PGK promoter. PGK promoter was amplified using primers WM17-1 TATCGAATTCTTCTACCGGGTAGGGGAGGCG and WM17-2 GACTGGATCCAGGTCGAAAGGCCCGGAGATGA and cloned between EcoRI and BamHI sites of pSMPP. pRK172 was a gift from Dr Michel Goedert, MRC Laboratory of Molecular Biology. NanoLuc-FRB-LgBiT and pNL1.1 were acquired from Promega.

### Cell lines

Cells were maintained at 37°C with 5% CO_2_ in a tissue culture incubator. All non-neuronal cells were cultured in complete medium composed of high-glucose DMEM (Gibco, 41966029) supplemented with 10% fetal bovine serum (Sigma, F4135) and 1% penicillin-streptomycin (Gibco, 15140122). LgBiT was sub-cloned from the NanoLuc-FRB-LgBiT control vector into the 2^nd^ generation pPMPP-lentiviral vector via PCR. Lentivirus was produced via co-transfection of pPMPP-LgBiT constructs, pcRV-Gagpol and pMD2G vectors into HEK-293T cells with lipofectamine 3000 according to manufacturer instructions (Thermo Fisher, LF300001). Target cells were infected with lentivirus supplemented with 5µg/ml polybrene (Sigma, TR-1003) and subsequently selected with puromycin 48 h post-infection.

### Protein Production and Purification

Protein purification was performed as previously described ^90^. Briefly, Human 6xHis-P301S-0N4R-tau-HiBiT was cloned into the bacterial expression vector pRK172 via PCR and expressed in BL21 DE3 competent *E. coli* (New England Biolabs, C2530H). Protein expression was induced with 500 μM IPTG overnight (12-16 h) at 16°C. Cells were pelleted and lysed in lysis buffer (1 mM benzamidine, 1 mM PMSF, 1X protease inhibitors, 14 mM β-mercaptoethanol, 300 mM NaCl, 25 mM HEPES, 30 mM imidazole, 1% NP-40). Lysates were cleared via ultracentrifugation and his-tagged protein purified on the AKTÄ Pure on the HisTrap HP column (GE Healthcare). Fractions of interest were second round purified via size exclusion chromatography on the Superdex 200 column (GE Healthcare). Fractions were pooled in PBS with 1mM DTT, concentrated to >3mg/ml and snap frozen in liquid nitrogen followed by long term storage at −80°C. To aggregate tau, 60µM tau monomer was incubated with 20 µM heparin, 2 mM DTT and 1X protease inhibitors in PBS for 24-72 h at 37°C shaking at 250RPM. Tau aggregation was quantified by THT fluorescence readout (excitation 440 nm; emission 510 nm) in the ClarioSTAR plate reader (BMG Labtech) with 5 µM tau aggregates and 15 µM THT.

### AAV Production and Titre

eGFP-P2A-LgBiT-nls was cloned into the pAAV-hSyn-eGFP vector genome plasmid via PCR to generate pAAV-hSyn-GPLN. Chimeric particles of adeno-associated virus produced with capsids of types 1 and 2 (AAV1/2) were prepared as previously described^91^. Briefly, pAAV-hSyn-GPLN, pAAV2/1, pAAV2/2 and pAdDeltaF6 helper plasmid were co-transfected in HEK293 cells. 48-60 h post-transfection, media and cells were harvested, lysed and viral particles subjected to iodixanol gradient ultracentrifugation (Sigma, D1556). The 40% iodixanol fraction was isolated and concentrated in PBS on Amicon 100 kDa spin columns and frozen at −80°C in single use aliquots. Genome titres were determined via qPCR targeting eGFP (Probe:5’-FAM/CCGACAAGC/ZEN/AGAAGAACGGCATCAA-3’) (IDTdna) and purity determined via SDS-PAGE followed by Coomassie staining.

### SDS-PAGE and Western Blotting

Samples for western blot were lysed in appropriate volumes of 1X RIPA buffer (Sigma, R0278) with protease inhibitors, lysates cleared via centrifugation and resuspended with appropriate volume of 4X NuPAGE LDS sample buffer (Thermo Fisher, NP0007) with 2 mM β-mercaptoethanol and boiled at 100°C for 10 minutes. Samples were subjected to SDS-PAGE using NuPAGE Bis-Tris 4-12% gels (Thermo Fisher, NP0324BOX) and electroblotted onto a 0.2 µm PVDF membrane using the Bio-Rad Transblot Turbo Transfer System. Transferred membranes were blocked in blocking buffer (5% milk in 0.1% tween 20 in 1X TBS (TBS-T)) for 1 h at room temperature, and incubated with primary antibody at desired concentration diluted in blocking buffer at 4°C overnight. Membranes were repeatedly washed with TBS-T and incubated with secondary HRP- or alexa-flour-conjugated antibody for 1 h at room temperature. Membranes were washed multiple times with TBS-T before being incubated with HRP substrate (Millipore, WBKLS0500) and membranes imaged using a ChemiDoc gel imager (Bio-Rad). Subcellular fractionation of the nucleus and cytosol prior to western blotting was performed via the REAP method, as previously described^92^

### Dissections and Primary Neuron Culture

All animal work was licensed under the UK Animals (Scientific Procedures) Act 1986 and approved by the Medical Research Council Animal Welfare and Ethical Review Body. Brains were removed from C57BL/6 mice and primary neurons were isolated as previously described^93^. The protocol was adapted to produce pooled hippocampal and cortical cultures. Post-dissection, 3×10^4^ cells were seeded per well into a white 96-well (Greiner bio-one, 655098) poly-L-lysine (RnD Systems, 3438-100-01) coated plate in plating medium composed of Neurobasal Plus (Gibco, A3582901) supplemented with 1mM GlutaMAX (Gibco, 35050061), 10% Horse Serum (Gibco, 26050070), 1% penicillin-streptomycin and 1x B-27 Plus (Gibco, A3582801) for 4 h before a complete media change to maintenance medium (plating medium devoid of serum).

### Human iPSC Maintenance, Culture and iNeuron Differentiation

Pluripotent iPSC cells were cultured on Vitronectin (Thermo Fisher, A14700) coated plates in feeder free conditions using complete TeSR™^-^E8 Basal medium (STEMCELL, 05991) and passaged using 0.5mM EDTA (Invitrogen, 15575-038). Once cells formed nice round cultures (approx. 1-2 weeks of culture), they were dissociated into single cells with accutase (Gibco, A11105-01) and plated on 0.1 mg/mL Poly-D-Lysine (Gibco, A38904-01) and Geltrex coated plates (Gibco, A14133-02)) at a density of 25,000 cells per cm^2^. Differentiation of iNeurons was performed as previously described ^69^. Differentiation was induced 24 h after plating (Day 0) in differentiation medium composed of DMEM/F12 (Gibco, 21331-020), 1 mM Glutamax, 1X non-essential amino acids (Gibco,11140-035), 1X N2 (Gibco, 17502001), 50 µM 2-mercaptoethanol (Gibco, 31350-010), 1X Antibiotic-Antimycotic (Gibco, 15240096) and 1 µg/ml doxycyline hyclate (Sigma, D9891-5G). From Day 2 the media was switched to maintenance medium composed of Neurobasal medium (Gibco, 21103-049), 1 mM Glutamax, 1X B27 (Gibco,12587-010), 10 ng/ml BDNF (Peprotech, 450-02), 10 ng/ml NT3 (Peprotech, 450-03), 1X Antibiotic-Antimycotic and 1µg/ml Doxycyline hyclate. On Day 4 the neurons were replated into 24 and 96 well plates at a density of 1*10^5^ cells per well and 5*10^4^ or 8*10^4^ cells per well respectively. Cells were infected with hSyn::AAV1/2-eGFP-P2A-LgBiT-nls particles at a cumulative multiplicity of 200,000 genome copies per cell on Day 5, 6 and 9. Media was replaced with fresh maintenance medium each day until Day 7 and then every 48 hours from then onwards. Tau entry assays, qPCR analysis, immunofluorescence and western blots were conducted on Day 14.

### iNeuron qPCR

Cells were collected on Day 0, Day 4, Day 7 and Day 14 for RNA extraction using Qiagen RNAeasy plus kit (74134). 350ng of RNA from each sample was converted into cDNA using Quantitect reverse transcription kit (Qiagen, 205313). Sample was diluted to 5 ng/µL and used in a 5 µL reaction with Luna universal qPCR master mix (New England biolabs, M3003L) and primers designed to detect a range of genes (**Fig. S6c**). qPCR was performed on the QuantStudio5 (Thermo Scientific). All samples were analysed in technical triplication from 3 independent differentiations. qPCR data was normalised to housekeeping gene 18s and results analysed using the ΔΔCt method.

### Genetic Knockdown

The following Human onTARGETplus SMARTpool siRNA were purchased from Horizon discovery: non-targeting pool (D-001810-10-05), RAB5A (L-004009-00-0005), RAB7A (L-010388-00-0005), VPS13D (L-021567-02-0005) and VPS35 (L-010894-00-0005). The following Mouse Accell SMARTpool siRNA were purchased from Horizon discovery: non-targeting pool (D-001910-10-05), Rab5a (E-040855-00-0005), Rab7a (E-040859-01-0005), Lrp1 (E-040764-00-0005), Vps13D (E-050935-00-0005), Npc1 (E-047897-00-0005) and Cyclophilin B (D-001920-20-05). Human onTARGETplus siRNA were diluted to 20 µM stock in RNase-free water and cells were transfected with 10 nM siRNA for 3 days with Lipofectamine RNAiMAX transfection reagent (Thermo Fisher, 13778075) in 6 well format. Knockdown cells were re-plated at day 3, and tau-HiBiT entry assayed at day 4. Mouse Accell SMARTpool siRNA were diluted to 100 µM in RNase-free water and added directly to neurons on DIV 4 at a final concentration of 1 µM. Tau entry assays were performed 72 h post-knockdown

### HEK 293T Tau Entry Assay

LgBiT expressing HEK 293T cells were seeded 2×10^4^ cells per well into white 96-well plates (Greiner bio-one, 655098) coated with poly-L-lysine (Sigma, P4707) in complete DMEM. 12-16 h later, the medium was replaced with 50 µl serum free CO_2_ independent medium (Thermo Fisher, 18045088) supplemented with 1 mM sodium pyruvate, 1% penicillin-streptomycin, 1 mM GlutaMAX and 50 µl tau-HiBiT solution added to final concentration in 100 µl. Cells were pre-treated for the indicated times with PitStop 2 (Sigma, SML1169), Dyngo 4a (Abcam, ab120689), dimethyl amiloride (Sigma, A125), heparin (Sigma, H3393), surfen hydrate (Sigma, S6951), or supplemented with lipofectamine 2000 (Thermo Fisher, 11668019). After tau incubation, media was aspirated, and cells washed once with PBS. PBS was aspirated and replaced with CO_2_ independent medium plus live cell substrate according to manufacturer instructions (Promega Nano-Glo Live Cell Assay System, N2013). The cells were incubated for 5 minutes and immediately loaded onto the ClarioSTAR plate reader where luminescent signal was quantified at 37°C. RLU data was normalised to viable cells per well acquired from a PrestoBlue viability assay when expressed as Cytosol entry(RLU/cell).

### Neuronal Tau Entry Assay

Primary neurons were infected at DIV 2 with hSyn::eGFP-P2A-LgBiT-nls AAV1/2 particles at a multiplicity of 50,000 genome copies per cell to express LgBiT. On day of assay, DIV 7 neurons were 100% media changed to fresh maintenance medium with desired concentration of tau-HiBiT in 100 µl. Neurons were pre-treated by freshly changing to medium supplemented with desired concentration of PitStop 2, Dyngo 4a, dimethyl amiloride, bafilomycin-A (Sigma, SML1661), methyl-β-cyclodextrin (Sigma, C4555), γ-cyclodextrin (Sigma, C4892), 25-hydroxycholesterol (Sigma, H1015) or heparin, followed by addition of tau-HiBiT to final desired concentration. Signal quantification was performed as described for the HEK293-T tau entry assay.

### PrestoBlue Viability Assay

Post-signal acquisition, PrestoBlue Cell Viability Reagent (Thermo Fisher, A13261) was added to cells according to manufacturer instructions and incubated for 42 minutes at 37°C and 5% CO_2_. The plate was loaded into the ClarioSTAR plate reader and fluorescence intensity quantified (excitation 540-570 nm; emission 580-610 nm). Total viable cells per well were calculated using a standard curve of known viable cells per well adjusted for background fluorescence.

### HEK 293T seeding assay

The seeding assay was performed as previously described^6^. Briefly, 20,000 cells were seeded into poly-L-lysine (Sigma, P4707) coated black 96-well plates in 50 µl OptiMEM (Thermo Fisher, 31985062). Tau assemblies were diluted in OptiMEM and 50 µl added to cells to final concentration in the presence or absence of LF. Cells were incubated at 37°C and 5% CO_2_ for 1 h before the addition of 100 µl complete medium to stop the transfection process. Cells were incubated at 37°C in an IncuCyte S3 Live-Cell Analysis System for 48–72 h after the addition of tau assemblies.

### Neuronal seeding assay

Neuronal seeding assays were performed as previously described^94^. Briefly, neurons were prepared as described and supplemented with a final concentration of 100 nM tau-HiBiT assemblies at 7 DIV in maintenance medium and incubated at 37°C and 5% CO_2_ until 14 DIV. At 14 DIV, cells were fixed with methanol, nuclei stained with DAPI and epitopes immunofluorescently labelled with anti-MAP2 and anti-mTau (T49) antibodies. Stained neurons were subjected to high-content imaging of 15 fields per well and tau aggregation quantified. mTau puncta per field were normalised to cell count per field. Any drugs were left on throughout the duration of the seeding assay.

### OHSC seeding assay

Slices were prepared from P301S tau transgenic pups aged 6-9 days as previously described^68^. Brains were extracted and maintained in slicing medium (EBSS + 25 mM HEPES) on ice. Brains were bisected along the midline and the medial surface attached to the stage of a Leica VT1200S Vibratome using cyanoacrylate (Loctite Superglue). Sagittal slices of 300 µm thickness were removed and the hippocampus was dissected using sterile needles. Slices were maintained on Millipore membranes (PICM0RG50) with 0.4 µm pore size in 6 well plates with 1 ml Slice Culture Medium as follows: 50% MEM with GlutaMAX (Gibco, 41090036) 18% EBSS, 6% EBSS + D-glucose, 1% penicillin-streptomycin, 0.06% nystatin (Gibco, 15340029) and 25% horse serum. Slices were maintained at 37°C and 5% CO_2_ in a humid atmosphere. Drugs were diluted in 1 ml culture medium and applied to the underside of the slices for 1 day before addition of 100 nM tau. 3 days later, media was removed and fresh medium added without drug or tau. 3 weeks post-challenge, OHSCs were fixed in 4% paraformaldehyde, nuclei stained with DAPI, and MAP2, phopsho-tau (AT8) and tau (DAKO) epitopes immunofluorescently labelled. Stained slice images were acquired using a Zeiss LSM780 confocal microscope with either a 20X or 63X objective lens and images stitched using ZEISS Zen software package.

### Plasmid-based endosomal lysis assay

HEK 293 cells were plated at 1×10^4^ cells per well in 96 well plates. The next day, media was exchanged for serum-free complete medium with pNL1.1 (Promega) plasmid at 10 ng/µl. Recombinant tau monomers or heparin-assembled tau were added to the media. Lipofectamine 2000 and endosome-destabilising adenovirus type 5 (ViraQuest) were added as positive controls for entry. Plates were examined for NanoLuc luminescence after 24 h on the ClarioSTAR plate reader.

### Transferrin uptake assay

5×10^5^ HEK 293T cells were suspended in serum-free complete medium and pre-treated with PitStop 2 (20 µM), Dyngo 4a (20 µM) or DMSO (0.1%) for 30 minutes at 37°C and 5% CO_2_. 50 µg/ml alexa-flour-647-conjugated human transferrin (Thermo Fisher, T23366) was added to the cell suspension and incubated on ice for 5 minutes, followed by 10 minutes at 37°C and 5% CO_2_. Cells were pelleted (500 x g, 5 minutes), acid washed twice (100 mM glycine, 150 mM NaCl pH 2.5 in PBS), fixed in 4% PFA (10 mins at room temperature) and subsequently resuspended in FACS buffer (0.5% bovine serum albumin in PBS). Transferrin uptake was then quantified via flow cytometry (CytoFLEX flow cytometer, Beckman Coulter).

### Live Flow Cytometry

Cells were rinsed once with 1X PBS and incubated with accutase for 12-15 minutes and allowed to detach. Cells were then resuspended in complete medium (HEK293) or maintenance medium (neurons/iNeurons) and flow cytometry performed in 96-well format on the Beckman Coulter CytoFLEX. 1*10^4^ events per condition were the minimum recorded events.

### Immunofluorescence

Media was aspirated and cells were rinsed once with PBS, then fixed with 4% PFA for 15 minutes at room temperature. Fixed cells were then rinsed 3 times with PBS followed by permeabilization with 0.1% triton X-100 (Sigma, X100) in PBS for 10 minutes at room temperature. Cells were then washed 3 times with PBS, and blocked with 1% BSA in PBS (IF block) for 1 h at room temperature. Primary antibody was diluted to required concentration in IF block and incubated overnight at 4°C. Antibody was removed, cells were then rinsed 3 times with PBS and incubated with alexa-flour conjugated secondary antibody in IF block for 1 h at room temp. Secondary antibody was removed, cells rinsed with PBS 3 times and imaged via fluorescence microscopy. Filipin III staining was performed according to manufacturer instructions (Sigma, SAE0087). For methanol fixation, cells were fixed and permeabilised with ice cold methanol on ice for 3 minutes, followed by 3 half washes with PBS (1X volume added, 0.5X total volume removed) followed by complete aspiration and 3 full washes with PBS. Permeabilised cells were then blocked and stained as described.

### Transmission Electron Microscopy

Formation of Tau-HiBiT fibrils was confirmed by uranyl acetate negative stain transmission electron microscopy at the Cambridge Advanced Imaging Centre as previously described^95^.

### Image analysis

Puncta corresponding to seeded aggregation in HEK 293T and primary neurons were quantified in Fiji using the ComDet plugin ^96^. Total puncta per field was normalised to cell count per field via DAPI staining or cell confluency detected by the IncuCyte S3 Live-Cell Analysis software. In neuronal seeding assays, only small puncta colocalised to neurons were quantified. AT8 staining in OHSCs was segmented into a binary threshold and fields of 150 µM x 150 µM were analysed for % AT8 positive area from slices from N=6 mice per condition.

### Statistical Analyses

All statistical analyses were performed via GraphPad Prism software. Differences between multiple means were tested by one-way ANOVA, followed by Tukey’s post hoc test unless otherwise indicated. Differences between two means were tested by unpaired t-test with Welch’s correction. All data represent mean values ± s.e.m with the following significances: ^ns^p > 0.05; **\***p ≤ 0.05; **\*\***p ≤ 0.01; **\*\*\***p ≤ 0.001; and **\*\*\*\***p ≤ 0.0001.

