## Supplementary Information for "Tau assemblies enter the cytosol in a cholesterol sensitive process essential to seeded aggregation"

### Figure S1

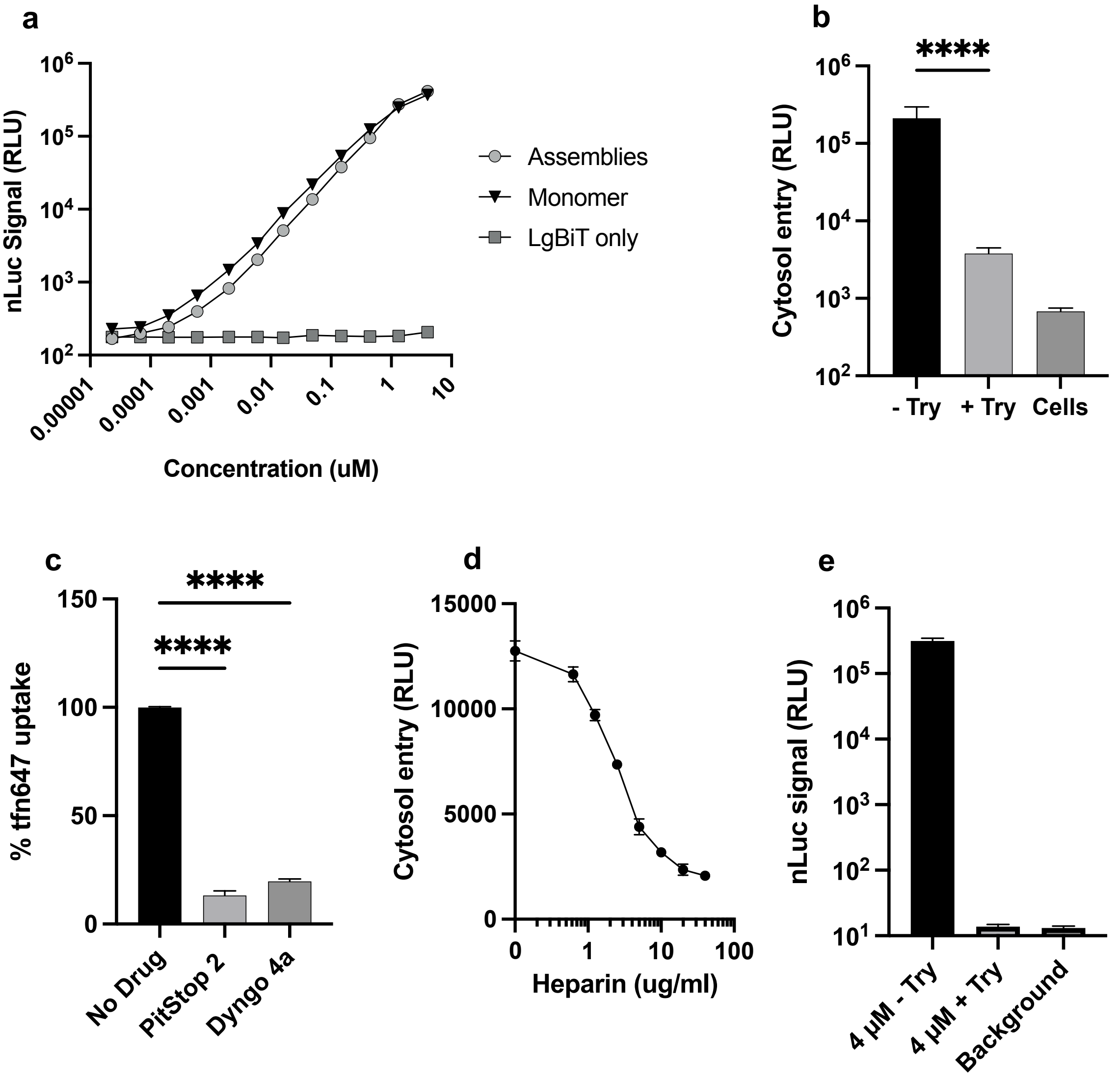


**Figure S1: a)** Representative titration curve of tau-HiBiT monomer or assemblies complexed with recombinant LgBiT. **b)** 10 minutes trypsin protease treatment of HEK-LgBiT treatment prior to signal acquisition after incubation with 250 nM tau-HiBiT assemblies for 1 h, n=6. **c)** In vitro (cell free) reconstitution of NanoLuc with 4 µM tau-HiBiT assemblies complexed with recombinant LgBiT in the presence or absence of trypsin. n=2-9 **d)** Quantification of alexa-647 labelled transferrin uptake assays in HEK293 cells in the presence of uptake inhibitors PitStop 2 (20µM) or Dyngo 4a (20 µM) via flow cytometry. $1*10^4$ events analysed per condition, N=3 independent biological experiments. **e)** Representative titration curve of NGL cells pre-treated for 30 minutes with increasing concentrations of heparin and entry of 50 nM tau-HiBiT assemblies assayed after 1 h, n=2 per data point. RLU; relative light units, Tfn; transferrin, Try; trypsin. Error bars are mean ± s.e.m.

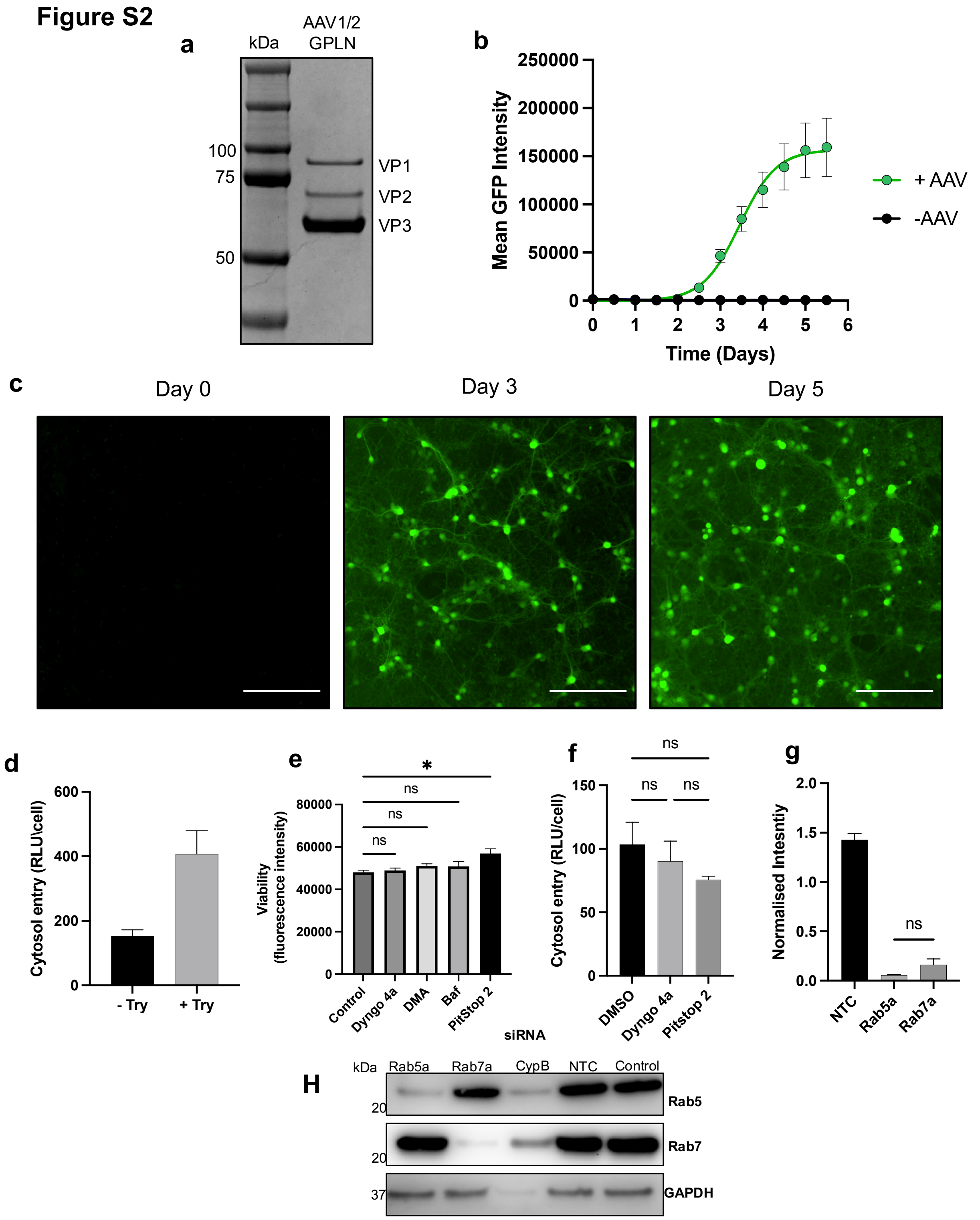


**Figure S2: a)** Coomassie staining of purified AAV1/2-hSyn-GPLN. **b)** Time course of GFP expression in DIV 2 WT neurons infected with AAV-GPLN at a multiplicity of 50,000 genome copies/cell over the course of 5 days; n=6, 5 fields per well analysed. **c)** Representative images of AAV transduced WT neurons from panel **b** at depicted time points. Scale bars, 200 µm. **d)** Treatment of DIV 7 GPLN-neurons with or without trypsin for ten minutes prior to signal acquisition after incubation with 50 nM tau-HiBiT assemblies for 1 h. In +try condition, we observed rounding of cells, likely amplifying luminescent signal. **e)** Representative presto blue viability assay of DIV 7 GPLN-neurons after tau entry assay and Dyngo 4a (40 µM), DMA (250 µM), Baf (200 nM), PitStop 2 (40 µM), or solvent control (0.1%, control) treatment throughout, showing no significant reduction in viability with any drug compared to control; n=3. **f)** Pre-treatment of DIV 14 GPLN-neurons with Dyngo 4a (20 µM), PitStop 2 (20 µM) or DMSO (0.1%) for 1 h followed by a 1 h entry assay with 50 nM tau-HiBiT assemblies. Neurons were transduced at DIV 9; n=3. **g)** Quantification of western blotting neuronal lysates 72 hours post-knockdown with Rab5 and Rab7 siRNA from three independent biological experiments. **h)** Full western blot from **fig 4m** (main text). Normalised intensity = pixel intensity of band /pixel intensity of corresponding GAPDH. Baf; bafilomycin-A, CypB; Cyclophilin B, NTC; non-target control, RLU; relative light units. Error bars are mean ± s.e.m.

**Figure S3**

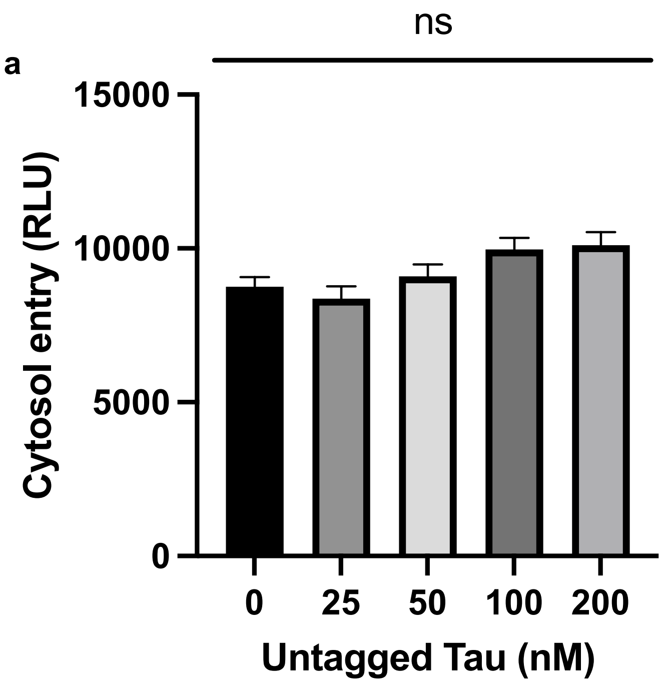


**Figure S3: a)** Titration of untagged tau assemblies in the presence of 50 nM tau-HiBiT assemblies in HEK-NGL cells and entry quantified after 1 h. n=4. Error bars are mean ± s.e.m.

**Figure S4**

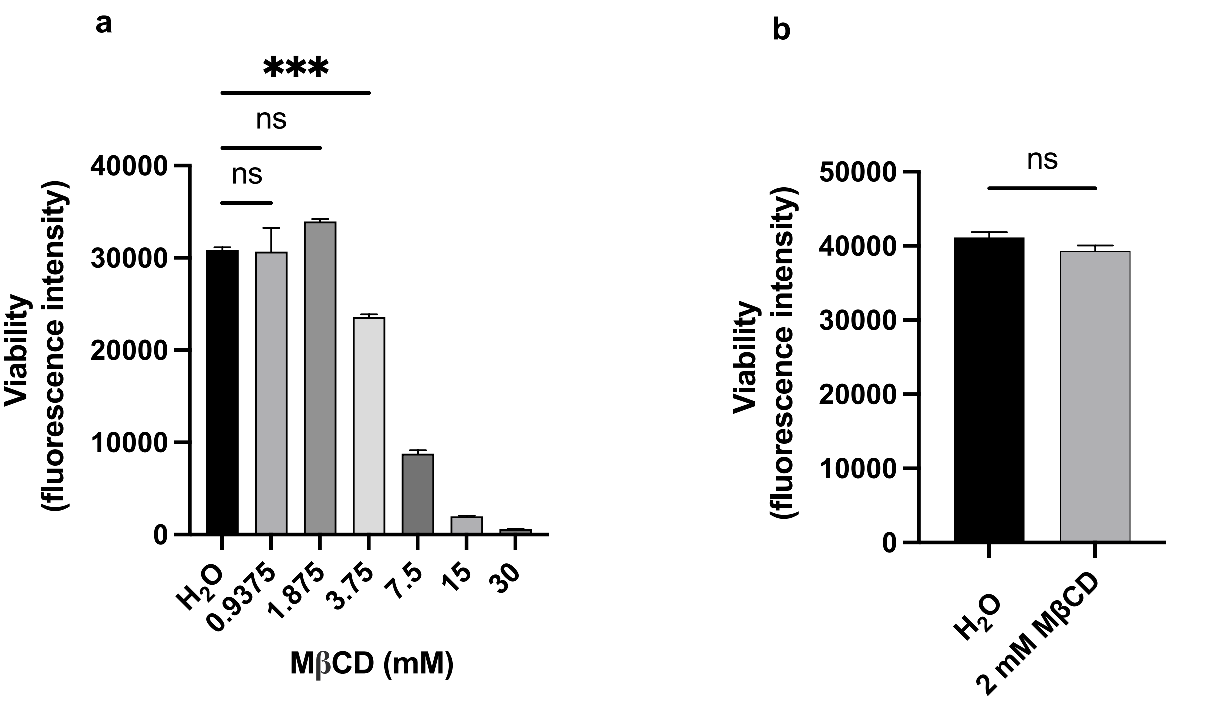


**Figure S4: a)** Presto blue viability assay of DIV 7 GPLN-neurons after 30 minutes pre-treatment with increasing concentrations of MβCD followed by a 1 h entry assay; n=3 **b)** Presto blue viability assay of DIV 7 GPLN-neurons after pre-treatment with 2 mM MβCD for 2 h followed by a 1 h entry assay. n=6. Error bars are mean ± s.e.m.

### Figure S5

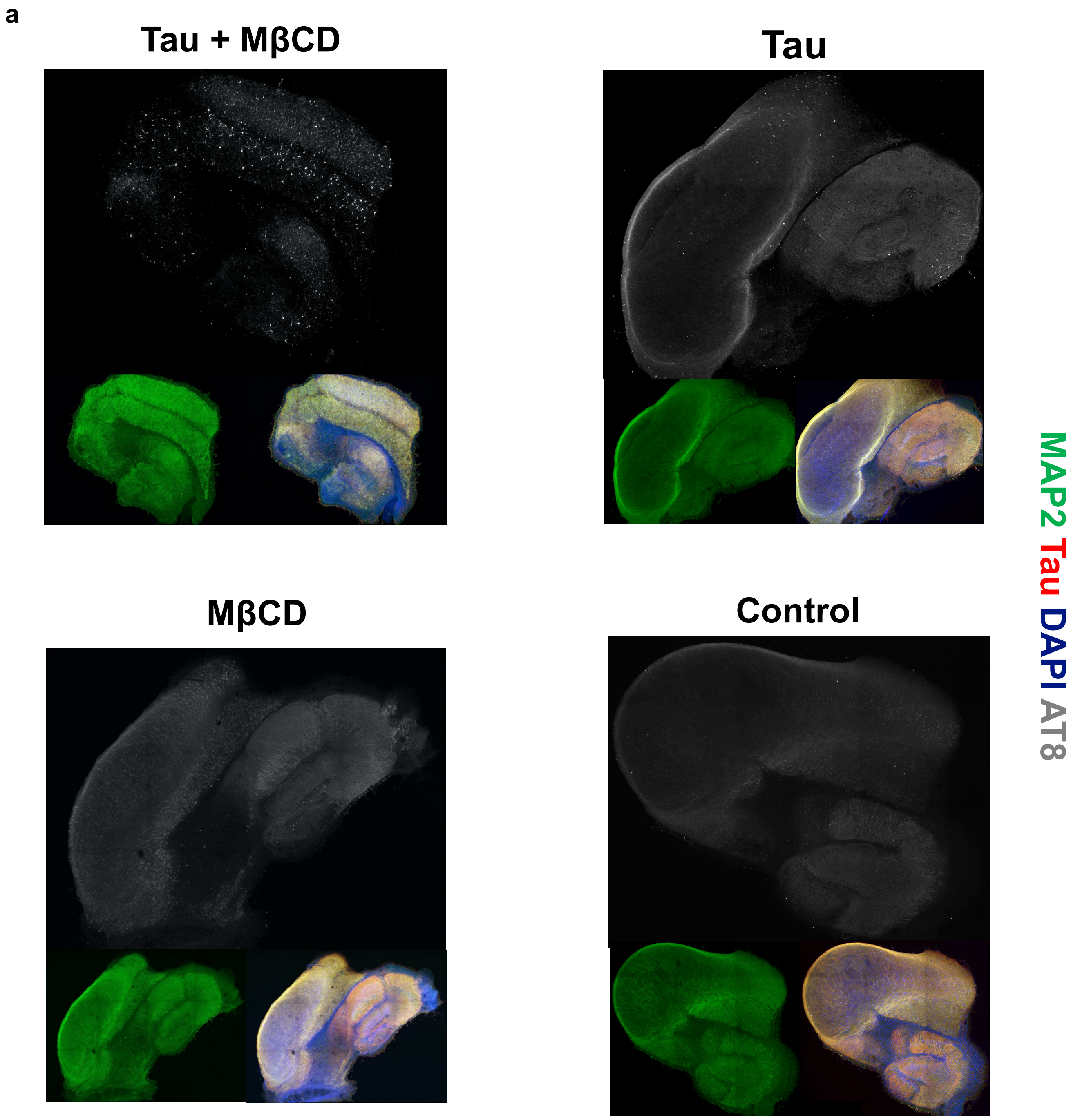


**Figure S5: a)** 10X magnification images of entire OHSC slices after a seeded aggregation assay treated with MβCD (200 μM) plus 100 nM tau assemblies, 100 nM tau only, MβCD (200 μM) only, or control solvent treated (0.1% EtOH).

### Figure S6

**a**

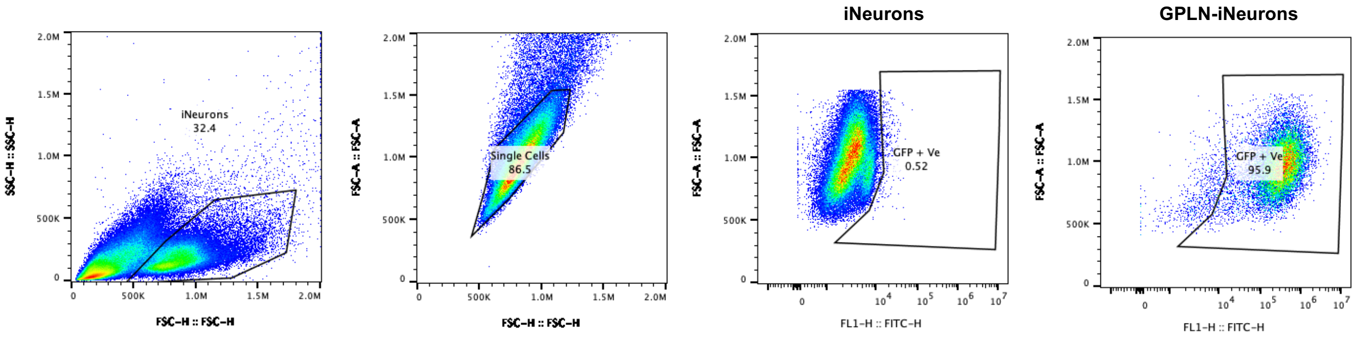


**b**

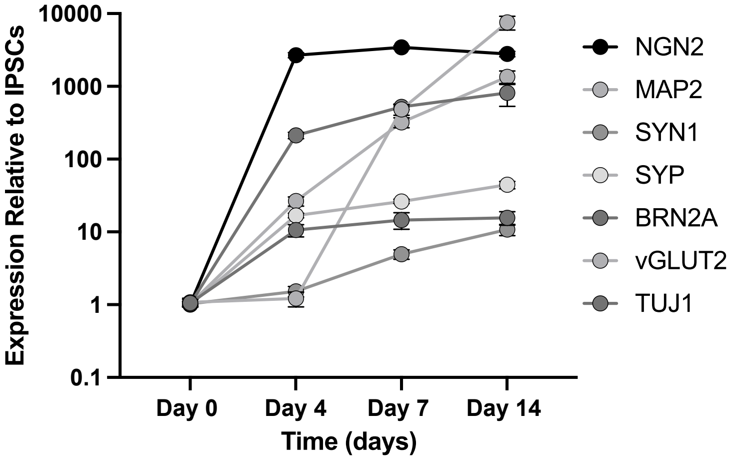


**c**

| Name | Sequence (5'-3') |
|---|---|
| **BRN2_fw** | ACCCGCTTTATCGAAGGCAA |
| **BRN2_rv** | CCTCCATAACCTCCCCCAGA |
| **VGLUT2_fw** | GTAGACTGGCAACCACCTCC |
| **VGLUT2_rv** | CCATTCCAAAGCTTCCGTAGAC |
| **TUBB3_fw** | CAACCAGATCGGGGCCAAGTT |
| **TUBB3_rv** | CCGAGTCGCCCACGTAGTT |
| **MAP2_fw** | AGACTGCAGCTCTGCCTTTAG |
| **MAP2_rv** | AGGCTGTAAGTAAATCTTCCTCC |
| **NGN2_fw** | TGTTCGTCAAATCCGAGACCT |
| **NGN2_rv** | CGATCCGAGCAGCACTAACA |
| **SYP_fw** | ACCTCGGGACTCAACACCTCGG |
| **SYP_rv** | GAACCACAGGTTGCCGACCCAG |
| **SYN1_fw** | CCCTGGGTGTTTGCCCAGAT |
| **SYN1_rv** | ACCACGGGGTACGTTGTACT |
| **NANOG_fw** | AGCAGATGCAAGAACTCTCCAA |
| **NANOG_rv** | TGAGGCCTTCTGCGTCACAC |
| **GRIA4_fw** | GGCCAGGGAATTGACATGGA |
| **GRIA4_rv** | AACCAACCTTTCTAGGTCCTGTG |
| **18s_fw** | GTAACCCGTTGAACCCCATT |
| **18s_rv** | CCATCCAATCGGTAGTAGCG |

**Figure S6: a)** Flow cytometry gating of day 14 iNeurons transduced with or without AAV1/2-hSyn-GPLN (GPLN-iNeurons). A minimum of $1*10^4$ events were recorded; n=6 per condition. **b)** qPCR of neuronal genes at day 0, day 4, day 7 and day 14 of the iPSC differentiation strategy to generate differentiated iNeurons; n=3 from N=3 independent differentiations. **c)** Primer sequences for genes targeted in qPCR. GPLN; eGFP-P2A-LgBiT-nls.
